# Efficient Disruption and Replacement of an Effector Gene in the Oomycete *Phytophthora sojae using CRISPR/Cas9*

**DOI:** 10.1101/025023

**Authors:** Yufeng Fang, Brett M Tyler

**Affiliations:** Interdisciplinary Ph.D. program in Genetics, Bioinformatics & Computational Biology, Virginia Tech, VA, USA; Center for Genome Research and Biocomputing and Department of Botany and Plant Pathology, Oregon State University, Corvallis, OR, USA

**Keywords:** genome-editing, effector, CRISPR/Cas9, Oomycetes, *Phytophthora sojae*

## Abstract

*Phytophthora sojae* is a pathogenic oomycete that infects soybean seedlings as well as stems and roots of established plants, costing growers $1–2 billion per year. Due to its economic importance, *P. sojae* has become a model for the study of oomycete genetics, physiology and pathology. Despite the availability of several genome sequences, the lack of efficient techniques for targeted mutagenesis and gene replacement have long hampered genetic studies of pathogenicity in *Phytophthora* species. Here, we describe a CRISPR/Cas9 system enabling rapid and efficient genome editing in *P. sojae*. Using the RXLR effector gene *Avr4/6* as target, we observed that in the absence of a homologous template, the repair of Cas9-induced double-strand breaks (DSBs) in *P. sojae* was mediated by non-homologous end joining (NHEJ), primarily resulting in short indels. Most mutants were homozygous, presumably due to gene conversion triggered by Cas9-mediated cleavage of non-mutant alleles. When donor DNA was present, homology directed repair (HDR) was observed, which resulted in the replacement of the target gene with the donor DNA. By testing the specific virulence of several NHEJ mutants and HDR -mediated gene replacements on soybeans, we have validated the contribution of Avr4/6 to recognition by soybean R gene loci, *Rps*4 and *Rps*6, but also uncovered additional contributions to resistance by these two loci. Our results establish a powerful tool for studying functional genomics in *Phytophthora*, which provides new avenues for better control of this pathogen.

## INTRODUCTION

*Phytophthora sojae* is a destructive pathogen which causes “damping off” of soybean seedlings as well as stem and root rot of established plants (Tyler, 2007). Morphologically and physiologically, oomycetes such as *P. sojae* resemble filamentous fungi, but evolutionally they are classified in the kingdom *Stramenopila*, which includes diatoms and brown algae (Tyler, 2001). Most *Phytophthora* species are plant pathogens that can damage a huge range of agriculturally and ornamentally important plants (Erwin & Ribeiro, 1996). For instance, *P. infestans*, which causes the potato late-blight disease, resulted in the Irish famine potato, and still is a problem for potato and tomato crops today (Judelson & Blanco, 2005). *P. sojae* causes around $1–2 billion in losses per year to the soybean crop (Tyler, 2007). Because of its economic impact, *P. sojae*, along with *P. infestans*, has been developed as a model species for the study of oomycete plant pathogens (Tyler, 2007).

The first two genome sequences of oomycetes (*P*. *sojae and P*. *ramorum*) were published approximately nine years ago (Tyler *et al.*, 2006), but functional genomics studies have been hampered by the lack of efficient strategies for genome engineering. DNA transformation procedures have been developed (Judelson *et al.*, 1993a, Judelson *et al.*, 1993b), but gene knockouts and gene replacements have never been possible because insertion of transgenes occurs exclusively by non-homologous end-joining (NHEJ) (Judelson, 1997, Tyler & Gijzen, 2014). Alternative approaches for functional analysis have included TILLING (Lamour *et al.*, 2006) and gene silencing (Judelson *et al.*, 1993b, Whisson *et al.*, 2005, Ah-Fong *et al.*, 2008, Wang *et al.*, 2011). However, TILLING, which is based on random mutagenesis, is very laborious and requires long term storage of large pools of mutants, and has not proven very useful in oomycetes. Gene silencing (RNAi), triggered using hairpin, antisense, and sense RNA constructs (Ah-Fong *et al.*, 2008) or using dsRNA directly (Whisson *et al.*, 2005, Wang *et al.*, 2011) has proven useful. However, knockdown of genes in oomycetes by RNAi is incomplete, and varies among gene targets, experiments and laboratories. Also, selective silencing of closely related genes is difficult.

Recent advances in engineered nucleases that specifically cleave genomic sequences in living cells have provided valuable tools to create targeted mutations in numerous organisms, from vertebrates, insects, and plants (reviewed in Gaj *et al.*, 2013) to microbes including parasites (Shen *et al.*, 2014, Wagner *et al.*, 2014, Peng *et al.*, 2015, Zhang *et al.*, 2014) and fungi (Jacobs *et al.*, 2014, Vyas *et al.*, 2015, Liu *et al.*, 2015). These nucleases, that include zinc finger nucleases (ZFN), transcription activator-like effector nucleases (TALEN), and CRISPR/Cas (Clustered Regularly Interspaced Short Palindromic Repeats / CRISPR associated), can generate a double-stranded break (DSB) at specific sites. By triggering repair of the DSB, either by error-prone NHEJ or homology directed repair (HDR), such methods can increase the rate of gene editing to levels that enable ready isolation of cells or organisms bearing a desired genetic change (Miller *et al.*, 2011). ZFNs and TALENs are engineered proteins containing a modular DNA recognition domain and a DNA cleavage domain. Owing to the complexity and limitations of selecting targets, ZFN has been gradually replaced by TALEN which has a simpler correspondence between its amino acid sequence and DNA recognition site (Miller *et al.*, 2011).

Like ZFNs and TALENs, the type II CRISPR/Cas9 system derived from the adaptive immune system of *Streptococcus pyogenes* also has DNA recognition and cleavage functions (Cong *et al.*, 2013, Mali *et al.*, 2013). However, DNA recognition is mediated by a single guide RNA (sgRNA) rather than a fused DNA recognition protein domain. The specificity of this system relies on the sgRNA which can direct the nuclease Cas9 to the target DNA sequence (Cong *et al.*, 2013, Mali *et al.*, 2013).

Here we have implemented the CRISPR/Cas9 system in *P. sojae*, using the RXLR effector gene *Avr4/6* (Dou *et al.*, 2010) as a target. RXLR effectors are a large superfamily of virulence proteins secreted by many oomycetes that have the ability to enter host cells in order promote host susceptibility (Jiang & Tyler, 2012). The presence of some RXLR effectors, such as Avr4/6, can be recognized by intracellular receptors encoded by plant resistance genes, triggering vigorous defense responses (Jiang & Tyler, 2012). The presence of Avr4/6 is recognized by soybean R genes *Rps*4 and *Rps*6 (Whisson *et al.*, 1994, Gijzen *et al.*, 1996, Dou *et al.*, 2010); recognition by *Rps*4 requires the N-terminus of Avr4/6, while recognition by *Rps*6 requires the C-terminus (Dou *et al.*, 2010). Our results demonstrate that CRISPR/Cas9-mediated gene disruption and gene replacement is an efficient and useful strategy for testing the function of specific genes in *P. sojae* such as *Avr4/6*, which should be useful for all oomycetes.

## RESULTS

### Establishment of the CRISPR/Cas9 system for *P. sojae*

To establish a CRISPR/Cas9 system for *P*. *sojae*, several functionalities had to be established, including efficient expression of Cas9 in *P. sojae* (our attempts to express TALENs in *P. sojae* were never successful due to strong gene silencing), efficient expression of guide RNAs (no oomycete RNA polymerase III promoters have been characterized), and targeting of the Cas9 enzyme to the *P. sojae* nucleus (commonly used mammalian nuclear localization signals function poorly in *P. sojae*) (Fang & Tyler, 2015).

For use in *P. sojae* we selected the *Streptococcus pyogenes* Cas9 encoded by a gene with human-optimized codons (*hSpCas9*), because this Cas9 version has been widely used in a variety of organisms (Cong *et al.*, 2013, Zhang *et al.*, 2014, Peng *et al.*, 2015), and matches *P. sojae* codon usage relatively well, with a codon adaptation index calculated at 0.776 (http://genomes.urv.es/CAIcal/E-CAI/). To test if this protein could be efficiently expressed in *P. sojae*, we fused GFP to the C-terminus of hSpCas9. Furthermore, to direct hSpCas9-GFP into the *P. sojae* nucleus, we used a strong synthetic NLS derived from a *P. sojae* bZIP transcription factor (Fang & Tyler, 2015), which we fused to the N-terminus of SpCas9 (Fig 1A). Expression of the NLS-hSpCas9-GFP construct in *P. sojae* transient and stable transformants resulted in a bright GFP signal strongly localized within the nuclei of *P. sojae* hyphae (Fig. 1A). These results indicated that hSpCas9 was strongly expressed in *P. sojae* without further codon optimization, and that the bZIP-derived NLS efficiently targeted the fusion protein to *P. sojae* nuclei.

**Fig. 1.**
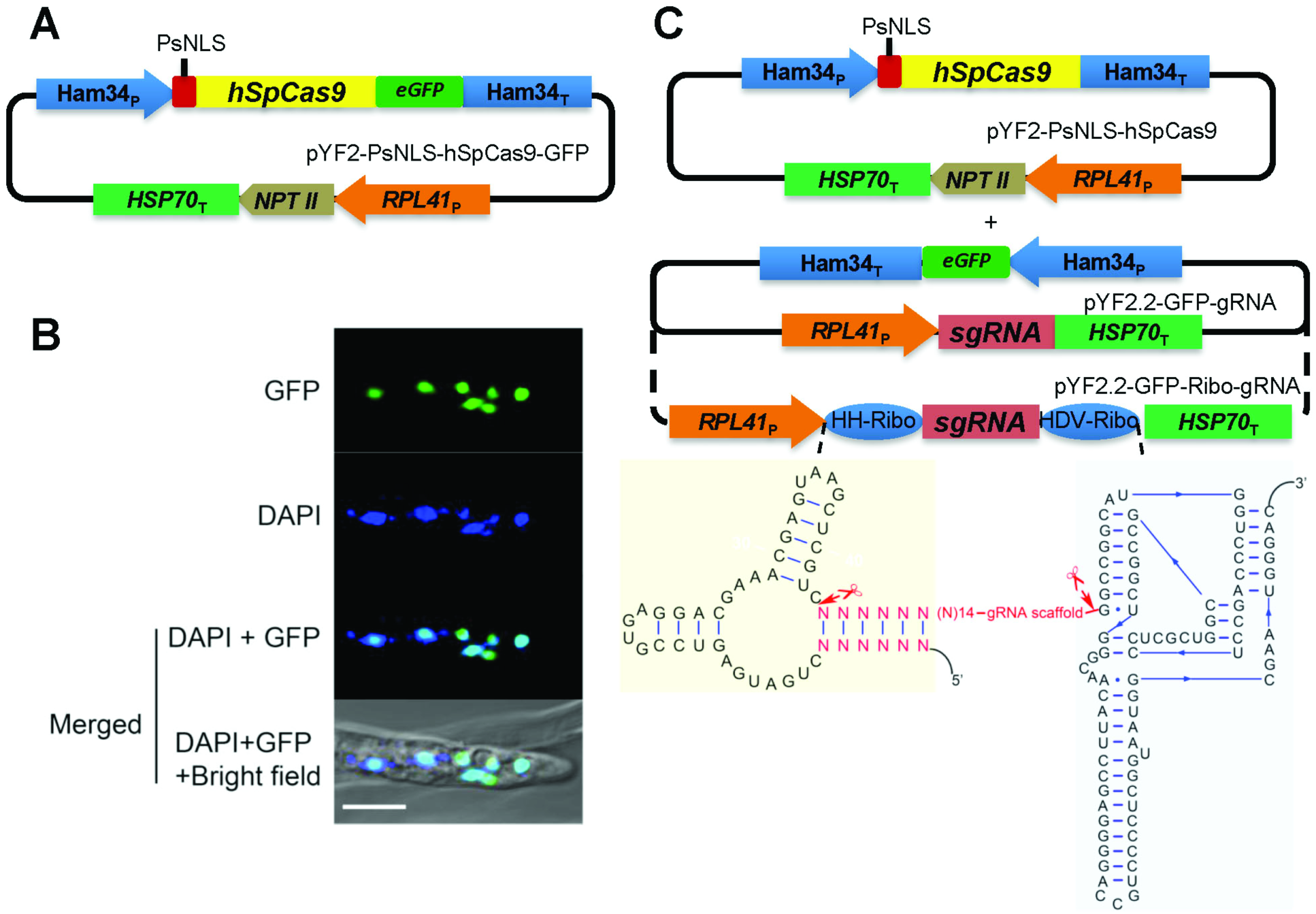
Cas9 and guide RNA constructs for *P. sojae* genome editing. (A) Top: Schematic of the plasmid for expression of *hSpCas9* fused to e*GFP* and an NLS in *P. sojae*. eGFP, enhanced green fluorescent protein. Bottom: *P. sojae* hyphae expressing PsNLS-hSpCas9-GFP from pYF2, counter-stained with DAPI; scale bars, 10 µm. (B) Top: Schematic of the plasmids used for expression of CRISPR constructs in *P. sojae.* Cas9 expression is driven by the Ham34 promoter, and cloned together with the selectable marker *NPTII* (encoding G418 resistance) driven by the RPL41 promoter (*P. sojae* ribosomal protein L41) in one plasmid. Transcription of sgRNA (including flanking ribozymes) is driven by the RPL41 promoter cloned into the same plasmid with an eGFP expression cassette (used as a screening marker). Bottom: Design of double ribozyme construct for release of sgRNAs from the primary RNA polymerase II transcript.

In most application systems, sgRNAs are synthesized by RNA polymerase III (RNA pol III), typically through use of a U6 small nuclear RNA (snRNA) promoter (Cong *et al.*, 2013, Mali *et al.*, 2013, Hwang *et al.*, 2013, Shen *et al.*, 2014, Zhang *et al.*, 2014). However, no RNA Polymerase III promoters have yet been functionally defined in oomycetes. To seek an oomycete U6 snRNA promoter, we aligned several U6 gene sequences found in *Phytophthora* and *H. arabidopsidis* genomes. The U6 transcripts were highly conserved among the different oomycetes (Fig. S1A), so we cloned the full length *P. sojae* and *P. infestans* U6 gene regions (Fig. S1B and C). To test the functions of the cloned U6 promoters, a 150bp fragment of the e*GFP* gene was inserted into each U6 coding region near the 3’ end (Fig. S1D). Surprisingly however, we did not detect any transcripts spanning the GFP reporter fragment by RT-PCR (data not shown). Therefore we sought an alternative approach for generating sgRNAs.

Recently, the generation of sgRNAs from RNA polymerase II promoters for genome editing was demonstrated in wheat (Upadhyay *et al.*, 2013), yeast (Gao & Zhao, 2014) and *Arabidopsis* (Gao *et al.*, 2015). The yeast and *Arabidopsis* systems used *cis*-acting ribozymes to trim flanking sequences from the sgRNAs, while the wheat system did not. Thus, we sought to use the constitutive *P. sojae RPL41* promoter (Dou *et al.*, 2008a, Dou *et al.*, 2008b) to direct the transcription of sgRNAs. Furthermore, two strategies were used for production of the sgRNAs. In the first, the sgRNA coding sequence was directly fused to the promoter without the addition of ribozyme sequences. In the second, the sgRNA coding sequences were flanked on the 5′ side by a hammerhead (HH) ribozyme and on the 3′ side by a HDV ribozyme (Gao & Zhao, 2014)(Fig. 1B).

To simplify the generation and screening of *P. sojae* transformants, the hSpCas9 gene and a resistance selection marker (*NPT II*) were placed in one plasmid, while the sgRNA gene together with a GFP marker gene were placed in a second plasmid (Fig. 1B). We also modified the *P. sojae* transformation procedure to enhance the efficiency of *P. sojae* transformation (see EXPERIMENTAL PROCEDURES).

### Cas9-mediated mutagenesis of *Avr4/6*

To test the activity of the sgRNA:Cas9 technology for *P*. *sojae* genome engineering, we selected as a target a *P. sojae* gene encoding an RXLR avirulence effector, Avr4/6 (GenBank: GU214064.1). *Avr4/6* is a single copy gene with no close paralogs. Furthermore, loss of *Avr4/6* function was expected to confer a phenotype that would not affect *in vitro* growth, namely the ability to successfully infect soybean cultivars containing resistance genes *Rps*4 or *Rps*6 (Dou *et al.*, 2010). sgRNAs targeting *Avr4/6* were designed based on the guidelines and web tool, *sgRNA Designer* (Doench *et al.*, 2014). Then sgRNA candidates rated highly by the web tool were further filtered by off-target analysis. Finally, two sgRNAs (sgRNA version A and B) were selected in which the respective Cas9 cleavage sites overlapped unique restriction enzymes sites (*Bst* UI and *Tsp* 45I respectively) that could be used to rapidly screen for mutations (Fig. 2A).

**Fig. 2.**
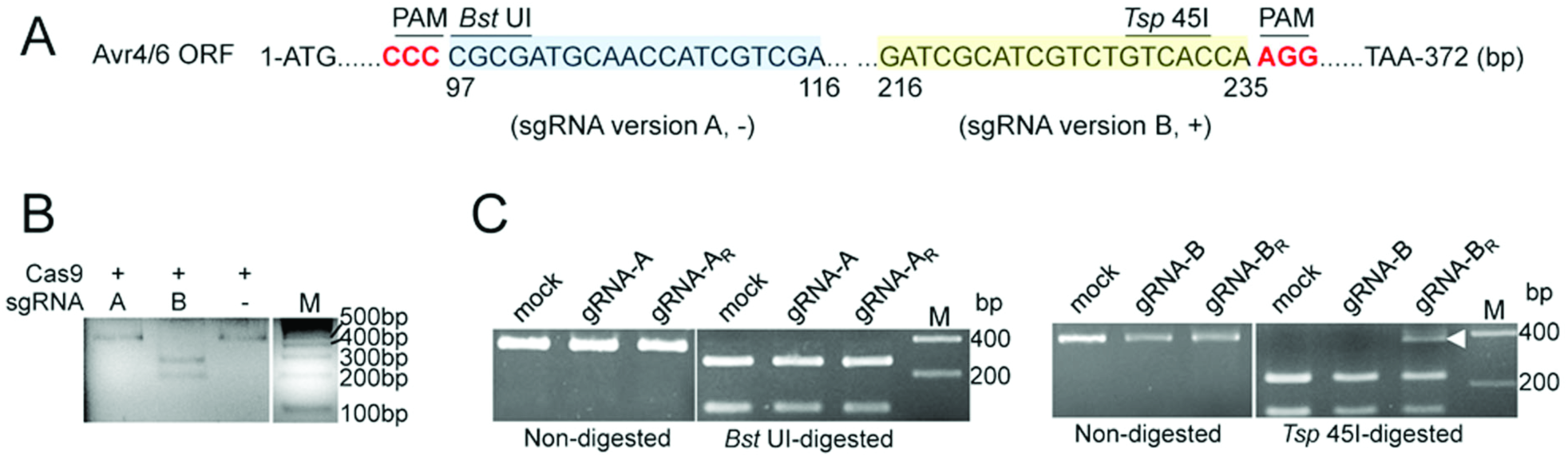
Design and evaluation of sgRNAs for targeting of *Avr4/6*. (A) Two sgRNA target sites within the *Avr4/6* ORF. Target sites of sgRNA A (sgRNA-A) and B (sgRNA-B) are highlighted in blue and yellow, respectively. sgRNAs A and B target *Avr4/6* on the negative (-) and positive DNA strand (+), respectively. The sgRNA-A site overlaps with a *Bst* UI restriction enzyme site (CGCG) and sgRNA-B with a *Tsp* 45I site (GT^G^/_C_AC). PAM, Protospacer Adjacent Motif is in bold red. (B) *In vitro* cleavage assay indicating Avr4/6 sgRNA-B can direct Cas9 cleavage of target PCR products but sgRNA-A cannot. DNA template was amplified from pBS-Avr4/6 by using M13F and M13R (Supplemental material). The colors of the original gel are inverted for clarity. (C) PCR and restriction enzyme analysis of *P. sojae* pooled transient expression transformants indicating that only sgRNA-B flanked by ribozymes (sgRNA- B_R_) produced amplicons resistant to restriction enzyme cleavage (arrowhead). Approximately 25-30% of the amplicon was resistant to *Tsp* 45I digestion. Experiment was performed in triplicate. sgRNA -A and -B, sgRNA lacking ribozymes; gRNA-A_R_, -B_R_, gRNAs flanked by ribozymes. Mock, *P. sojae* transformants only receiving Cas9 plasmid. In (B) and (C), all gel panels placed together were from the same gel; white dividers indicate lanes not adjacent in those gels.

To examine the activity of the designed sgRNAs, Avr4/6-A and Avr4/6-B, we carried out a sgRNA-mediated *in vitro* cleavage assay of target DNA. In these assays, the sgRNAs were synthesized using T7 RNA polymerase, and SpCas9 protein was purchased from New England Biolabs. Cas9/sgRNA-B completely cleaved the target DNA, whereas Cas9/sgRNA-A showed no activity *in vitro* (Fig. 2B).

In parallel with the *in vitro* assays, we used transient expression in *P. sojae* protoplasts to test whether the hSpCas9 and sgRNAs produced *in vivo* could modify the endogenous *Avr4/6* gene. The two *Avr4/6*-specific sgRNAs were assembled into the *P. sojae* expression plasmid under the control of the *RPL41* promoter, either flanked with ribozymes (Avr4/6-sgRNA-A_R_, -B_R_) or without ribozymes (Avr4/6-sgRNA-A, -B). The sgRNA constructs were co-transformed with the hSpCas9 expression plasmid into *P. sojae* strain P6497 by PEG-mediated protoplast transformation. Transformants were enriched by G418 selection 12 h after transformation when hyphae had regenerated. After 24 h, DNA was extracted from the culture containing the pooled transformants. *Avr4/6* sequences were amplified from the pool of genomic DNAs and screened for mutants resistant to the relevant restriction enzymes (*Bst* UI for A and *Tsp* 45I for B). We found that the *Avr4/6* amplicons from the two sgRNA-A transformations (constructs with and without ribozymes) were still fully subject to restriction enzyme cleavage, indicating failure of the Cas9-mediated mutagenesis (Fig. 2C). In contrast, the transformation utilizing the sgRNA version B flanked by ribozymes yielded restriction enzyme resistant amplicons (Fig. 2C). However the transformation utilizing the sgRNA version B without ribozymes did not yield restriction enzyme resistant amplicons (Fig 2C). To validate the enzyme cleavage results, we further sequenced the nested PCR products amplified from the enzyme digestion products from the sgRNA-B_R_ transformants. The sequence chromatograms showed pure sequences proximal to the target site, and mixed sequences distal to the target site (data not shown), suggesting the presence of indel mutations at the target site. These observations indicated that in *P. sojae*, RNA polymerase II can be successfully used for generating sgRNA, provided that ribozymes are employed to remove the surrounding sequences from the transcripts. In addition, based on the intensity of the restriction enzyme-resistant band on the agarose gel relative to the intensity of the fragments resulting from *Tsp* 45I cleavage, around 25-30% of the amplicons appeared to carry a mutation in the restriction site (Fig. 2C). The failure of the sgRNA version A may result from strong self-complementarity that we subsequently discovered in its sequence using *RNA structure (http://rna.urmc.rochester.edu/RNAstructureWeb/Servers/Predict1/Predict1.html)*, which could block its binding to target DNA.

To characterize CRISPR/Cas9-generated *Avr4/6* mutations in detail, the transformation with the ribozyme-containing Avr4/6-sgRNA-B_R_ construct was repeated. Individual G418-resistant transformants were isolated and screened for the presence of GFP indicating the presence of the sgRNA construct. Of 50 primary transformants screened, 6 exhibited green fluorescence. Of these, 4 yielded *Avr4/6* amplicons that were partially or fully resistant to *Tsp* 45I digestion (Fig. 3A), indicating the presence of *Avr4/6* mutations. Since *P. sojae* protoplasts and hyphae are multinucleate, and hence might be expected to harbor nuclei with a diversity of *Avr4/6* mutations, we isolated zoospores (which are mononucleate) from three of the primary transformants (T11, T18 and T32; the fourth, T30, did not produce zoospores). Ten single zoospore lines were isolated from each transformant, and the *Avr4/6* amplicons were screened by restriction enzyme digestion and by Sanger sequencing. T11, T18 and T32 yielded 7, 10 and 10 pure mutant lines respectively (summarized in Table 1). All 27 pure lines showed a homogeneous sequence profile (Fig. S2), indicating that all of them were already homozygous, and carried the same mutation in both alleles. All 10 of the T32 lines were homozygous for the same mutation. The lines derived from T18 included 9 lines homozygous for one Avr4/6 mutation and one homozygous for a different mutation. The lines derived from T11 included two homozygous for one mutation (mut1) and five homozygous for a second mutation (mut2). The three remaining lines were heterozygous and biallelic, containing a third mutation paired with mut1. (Fig. 3B and Fig. S2). No lines retained any wild type alleles, either homozygous or heterozygous.

**Fig. 3.**
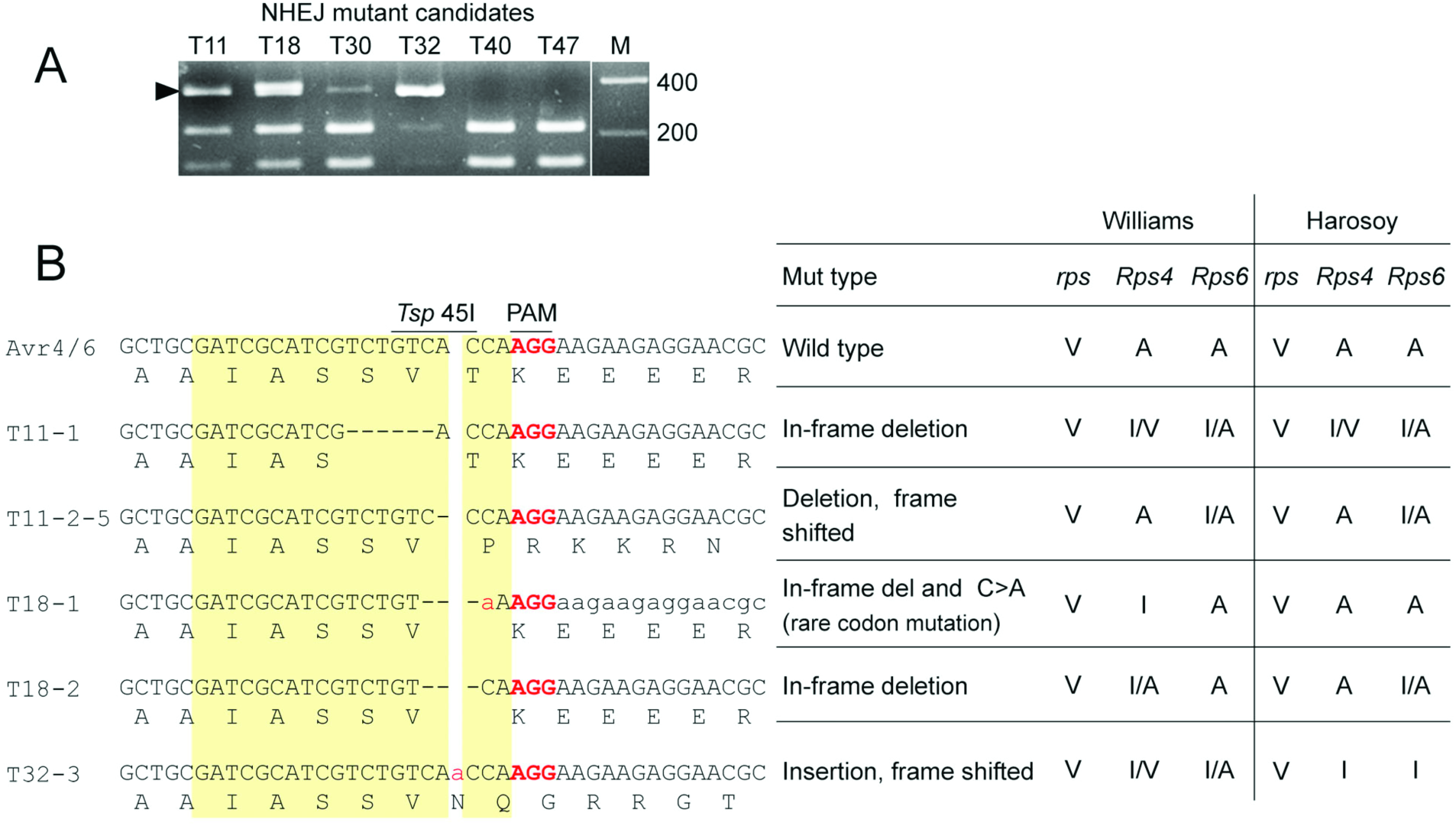
Characterization of individual *P. sojae* NHEJ-mediated mutants. (A) *Tsp* 45 I screening of Avr4/6 amplicons from six individual *P. sojae* transformants carrying *hSpCas9* and sgRNA-B_R_ plasmids. Arrowhead indicates *Tsp* 45 I-resistant amplicons. Transformants are expected to carry a mixture of modified and non-modified *Avr4/6* genes. (B) Left: Sequences of Avr4/6 mutant amplicons from single zoospore lines derived from transformants T11, T18 and T32 (T30 produced no zoospores). Target sites are highlighted in yellow and the PAM sequences are in bold red. (Right) Summary of virulence assays of Avr4/6 mutants on *Rps*4- and *Rps*6-containing cultivars. V, virulent; A, avirulent; I, intermediate; I/A intermediate to avirulent; I/V intermediate to virulent.

**Table 1.** ***Avr4/6* CRISPR/Cas-induced NHEJ mutations**

| Mutants | WT/WT<sup>a</sup> | Mutant patterns |  |  |  | Biallele<br>mut1/mut2,3 |
| --- | --- | --- | --- | --- | --- | --- |
|  |  | Homozygote<br>mut1/mut1 | mut2/mut2 | Heterozygote<br>WT/mut1 | WT/mut2 |  |
| T11<sup>b</sup> | 0 | 2 | 5 | 0 | 0 | 3* |
| T18<sup>c</sup> | 0 | 9 | 1 | 0 | 0 | 0 |
| T32<sup>d</sup> | 0 | 10 | 0 | 0 | 0 | 0 |

<sup>a</sup> WT, wild-type sequence
<sup>b</sup> T11, mut1, six nucleotides deletion; mut2, one nucleotide insertion; \*heterozygous mut1/mut3; mut3 is a three bp deletion. The sequencing profiles of the heterozygotes were disambiguated using the web-tool TIDE (Brinkman *et al.*, 2014)
<sup>c</sup> T18, mut1, three bp deletion; mut2 three bp deletion plus one bp substitution
<sup>d</sup> T32, mut1, one bp insertion;

Each of the mutations consisted of a short indel, located specifically at the Cas9 cleavage site, i.e. between the 17^th^ and 18^th^ nucleotide of the sgRNA target site. Deletions of one, three and six bp were observed, one contained a one bp insertion, and one combined a three bp deletion with a one bp replacement (Figure 3B and Table 1).

We also tested the inherent stability of the mutants by sub-culturing each of the single zoospore lines for at least three generations on media without G418 selection. All of the mutated sites examined remained the same as the first generation, based on the restriction enzyme cleavage assay and sequencing of the *Avr4/6* amplicon. In addition, we noticed that a few of the sub-cultured mutants had lost the ability to grow under G418 selection, showing that mutations were stable regardless of presumptive continued presence of the sgRNA:Cas9 construct. Interestingly, one transformant T47 which did not show obvious mutations in the first generation acquired the same single adenine insertion as T32-3 after sub-culturing of the unpurified transformant for one generation (Fig. S3), presumably because the sgRNA:Cas9 constructs were integrated into the genome and continued to actively cleave the target in each generation. Collectively, these results indicate that our CRISPR/Cas9 system can efficiently and specifically trigger the introduction of NHEJ mutations, typically short indels, into the *P. sojae* genome.

### Homologous gene replacement stimulated by the CRISPR/Cas9 system

Donor DNA-mediated repair of sgRNA-guided Cas9 cuts has proven an efficient way to facilitate gene replacements via homology-directed repair (HDR) (Cong *et al.*, 2013, Mali *et al.*, 2013). To determine if sgRNA:Cas9 mediated DSB could stimulate homologous recombination in *P. sojae*, we co-transformed the CRISPR constructs that were successfully used for mutation of *Avr4/6*, along with uncut donor DNA plasmids that contained the entire *NPT II* ORF flanked by different lengths of the sequences surrounding the *Avr4/6* gene. An equimolar ratio of the three plasmids was used (Fig. 4A). Since preliminary experiments had shown that expression of the *NPT II* gene from the *Avr4/6* promoter was insufficient for G418 selection, the *NPT II* gene was included in the Cas9 plasmid for selection of transformants. We used homology arms consisting of three different lengths of 5′ and 3′ flanking sequences, namely, 250 bp, 500 bp, and 1 kb, to assess which would enable the highest recombination efficiency (Fig. 4B). The *NPT II* gene in the Cas9 expression plasmid served as a negative control, because it lacked any *Avr4/6* flanking sequences.

**Fig. 4.**
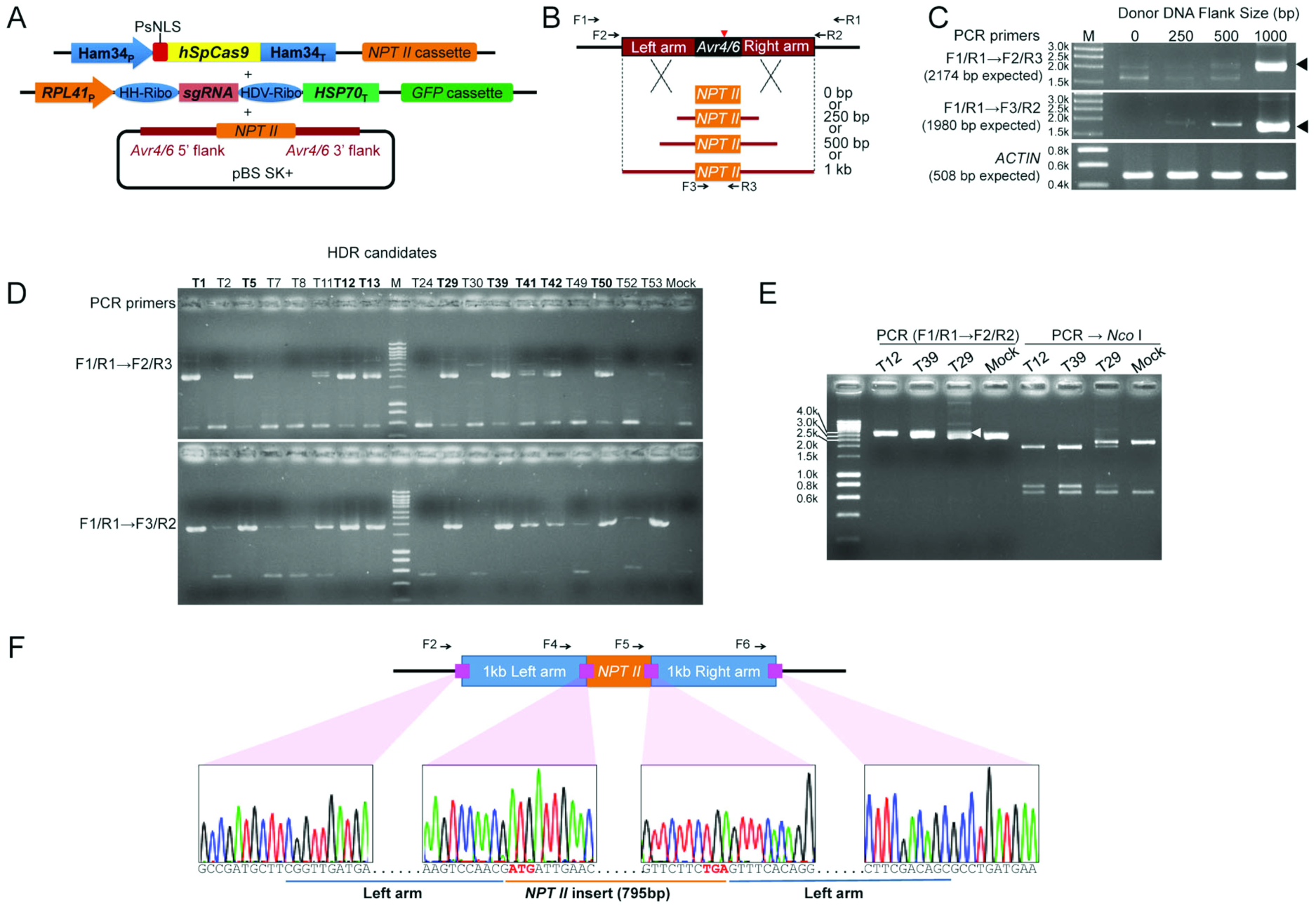
HDR-mediated replacement of the *Avr4/6* ORF with an *NPT II* ORF. (A) Strategy used for gene replacement. Plasmids containing a homologous donor DNA (*NPT II* with *Avr4/6* flanking sequences) were co-transformed with the Avr4/6 sgRNA-B_R_ and hSpCas9 constructs. (B) Three different sizes of homology arms, 250 bp, 500 bp and 1 kb, flanking the *Avr4/6* locus were used. The *NPT II* gene in the hSpCas9 expression plasmid served as the control (0 bp homologous arm, Mock). Primers used to screen the HDR mutants and validate the replaced region are shown as arrows. Primer pairs F1/R1, and nested primer pairs F2/R3, F3/R2 and F2/R2 were used for HDR mutant screening. Red arrow indicates the CRISPR/Cas9 cleavage site (between 232 bp and 233 bp of *Avr4/6* ORF). (C) Analysis of genomic DNA from pooled transformants produced using the four sizes of flanking sequences, using nested PCR. Arrowheads indicate sizes expected if HDR has occurred. *ACTIN* = actin control for DNA quality. (D) Screening of individual HDR transformants generated with the 1 kb flanking sequence plasmid. The 9 positive HDR mutants are highlighted in bold. (E) PCR analysis of representative zoospore-purified lines of HDR mutants, demonstrating that T12 and T39 are homozygotes while T29 is a heterozygote. DNA sizes (bp) before restriction enzyme cleavage: WT, 2936; HDR mutants, 3327; after *Nco* I digestion: WT, 2273 + 663; HDR mutant, 1945 + 751 +631. The arrowhead indicates the fragment amplified from the *NPT II*-replaced allele. Primer F4= Avr4/6_up500bp_PhusF (Table S1). (F) Sanger sequencing traces of junction regions confirming that the *Avr4/6* ORF was cleanly replaced by the *NPT II* ORF in a representative zoospore purified clone (HDR-T12-1). Start and stop codons are in red bold.

Following co-transformation and G418 enrichment, the bulk transformants were subjected to genomic DNA extraction and PCR analysis. PCR amplifications using primers located outside the *Avr4/6* homology arms and within the *NPT II* gene were used to detect homologous recombination events. The results suggested that HDR had occurred, but the frequency was variable depending on the length of the flanking sequences in the donor DNA plasmids. The transformant population generated with the 1 kb flanking sequences showed the highest frequency of gene replacement. The population generated with the 500 bp flanking sequences showed a much lower frequency compared to the 1 kb population. The population from the 250 bp flanking sequences showed very low recombination frequencies (Fig. 4C).

Next, to characterize HDR events in detail, we generated single *P. sojae* transformants derived from the 1 kb arm donor. After screening 68 individual G418 resistant transformants for GFP production, we identified 18 transformants bearing the two CRISPR components. Then, using nested primers specific for HDR events (Fig. 4B), we found evidence for gene replacement events in 9 of the transformants (Fig. 4D). Sanger sequencing across the junctions of the flanking sequences and *NPT II* in the nested PCR products was also consistent with replacement of the *Avr4/6* ORF with *NPT II* gene (data not shown). Three HDR mutants, namely HDR -T12, -T29 and -T39, that readily produced zoospores, were selected for functional tests. After zoospore isolation, the *Avr4/6* region of each single zoospore line was examined by PCR amplification using primers flanking the two homologous arms and cleavage of the amplicon by the restriction enzyme *Nco* I. We found that all of the 11 single zoospore lines obtained from HDR-T12, and all 8 obtained from HDR-T39 were homozygotes (Fig. 4E); this was further validated by Sanger sequencing (Fig. 4F). In contrast, all 20 of the single zoospore lines of T29 appeared to be heterozygotes that contained a HDR event in just one of the two *Avr4/6* alleles. This was further verified by PCR amplification using primers outside of homology arms and in *NPT II* gene. More detailed analysis of three of the HDR-T29 lines revealed that the non-HDR alleles possessed the same mutation, an adenine deletion, in every case (Fig. S3), presumably caused by NHEJ.

### Modified recognition of *Avr4/6* mutants by soybeans carrying the *Rps*4 and *Rps*6 loci

In order to test the effects of the CRISPR/Cas9-induced *Avr4/6* mutations on *P. sojae* recognition by plants containing the *Rps*4 and *Rps*6 loci, the five homozygous NHEJ mutants and two homozygous HDR mutants were inoculated onto hypocotyls of soybean isolines containing *Rps*4 (L85-2352) or *Rps*6 (L89-1581) in a Williams background, as well as isolines containing *Rps*4 (HARO4272) or *Rps*6 (HARO6272) in a Harosoy background. 4 days after inoculation (dpi), the specific virulence of the different mutants was scored and analyzed by Fisher’s exact test (Table 2). We observed that the frameshifted mutants T32-3 and T11-2-5 both showed increased killing of *Rps*4- or *Rps*6-containing soybean seedlings in both Williams and Harosoy backgrounds (Table 2). The increased killing of both *Rps*4 and *Rps*6 plants by T32-3 was statistically significant (p < 0.01), while the increased killing by T11-2-5 of *Rps*4 but not *Rps*6 plants was significant (p<0.05). On the other hand, the increased killing in every case was still significantly (p < 0.05) less than the killing of *rps* plants lacking *Rps*4 or *Rps*6 (Table 2). Thus T32-3 was scored as intermediate on *Rps*4 and *Rps*6 plants, while T11-2-5 was scored as avirulent and intermediate respectively. The other NHEJ mutants, containing in-frame deletions, also showed increased killing of *Rps*4- and *Rps*6-containing plants, but significantly (p < 0.03) less killing than observed with *rps* plants. The *Avr4/6* mutant having a two amino acid deletion (T11-1) showed an intermediate phenotype that was close to fully virulent on *Rps*4 plants while the two mutants a single amino acid deletion (18-1 and 18-2) showed intermediate to avirulent phenotypes (Fig. 5; Table 2).

**Fig. 5.**
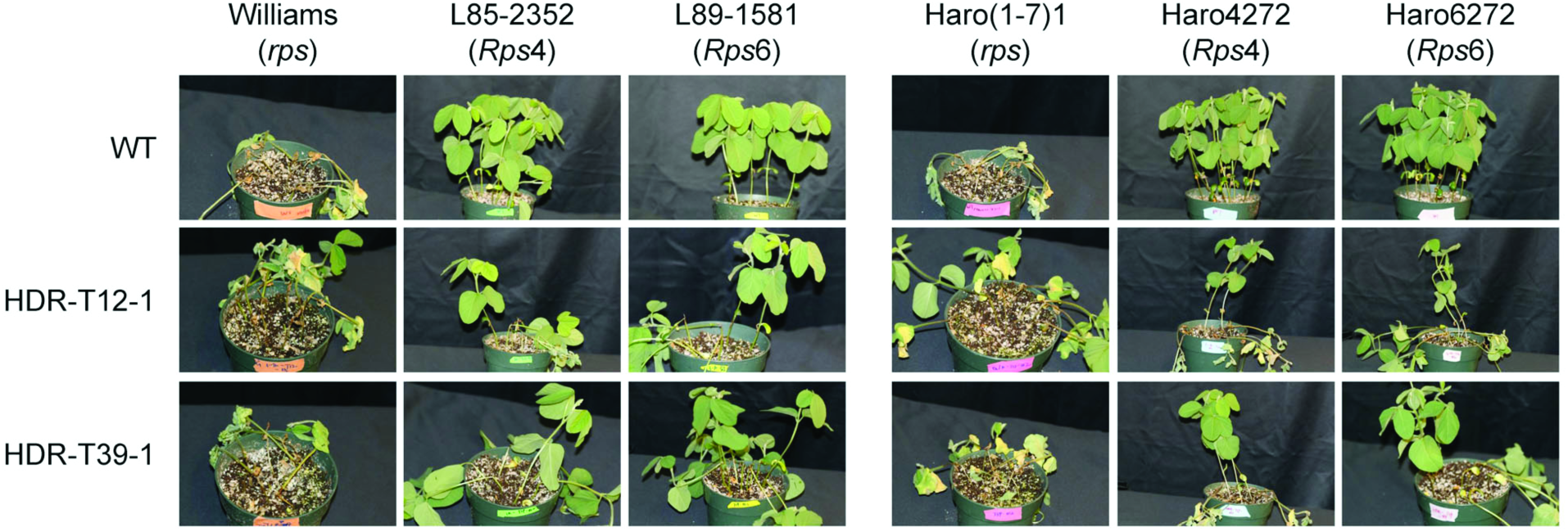
Infection phenotypes of Avr4/6 mutants Representative photos showing soybean seedlings inoculated on the hypocotyls with *P. sojae Avr4/6* HDR mutants. L85-2352 and HARO4272 contain *Rps4* while L89-1581 and HARO6272 contain *Rps6.* Photographs were taken 4 days post-inoculation.

**Table 2.**
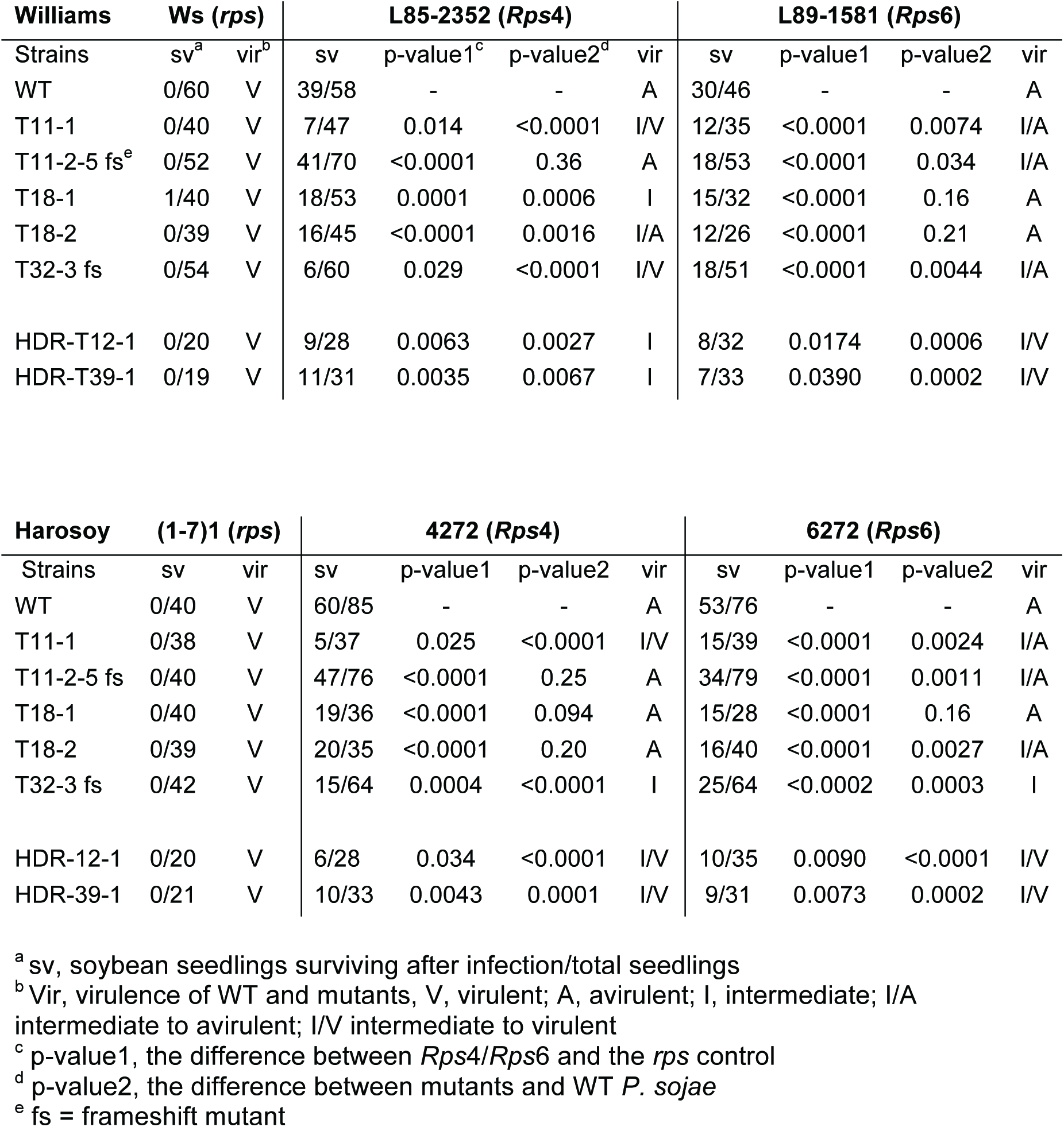
**Characterization of the virulence of *P. sojae Avr4/6* NHEJ- and HDR-mutants on soybeans**

The two homozygous HDR mutants (T12-1 and T39-1), both showed significantly (p < 0.01) more killing of *Rps*4- or *Rps*6-containing soybean seedlings in both Williams and Harosoy backgrounds (Table 2 and Fig.5), but the killing was significantly (p < 0.05) less than on *rps* plants. Thus, both mutants were scored as intermediate to virulent on *Rps*4 and *Rps*6.

## DISCUSSION

Substantial numbers of oomycete genomes have now been sequenced, and even larger numbers are underway (Jiang *et al.*, 2013, Jiang & Tyler, 2012, Kamoun *et al.*, 2014). These genomes contain 15,000 to 25,000 genes, of which approximately half in most species show the rapid sequence divergence expected of infection-related genes (Tyler *et al.*, 2006, Haas *et al.*, 2009, Baxter *et al.*, 2010, Jiang & Tyler, 2012, Jiang *et al.*, 2013). Many of the diverging genes encode large families of effectors, toxins, hydrolytic enzymes, hydrolytic enzyme inhibitors, and ABC transporters, as well as numerous proteins of unknown function (Tyler *et al.*, 2006, Haas *et al.*, 2009, Jiang & Tyler, 2012). To date however, the tools available for assessing the functions of these genes have been limited to RNAi-mediated gene silencing and to expression of ectopic transgenes. RNAi-mediated gene silencing has proven useful (Judelson *et al.*, 1993b, Ah-Fong *et al.*, 2008), especially with the introduction of transient silencing (Whisson *et al.*, 2005, Wang *et al.*, 2011). However the level of silencing is unpredictable and often unsatisfactory, and some genes have required a major effort to find a few silenced lines. Expression of ectopic transgenes has also proven useful. However major limitations have been the unpredictability of the expression level and the related issue of potential over-expression artifacts. The inability to eliminate the contribution of the endogenous gene has also limited the usefulness of ectopic transgenes.

Here we have adapted the CRIPSR/Cas9 sequence-specific nuclease technology for use in *Phytophthora sojae.* This involved overcoming several technical hurdles. One of these was the fact that commonly used mammalian nuclear localization sequences (NLS) do not function in *P. sojae* and that *P. sojae* nuclear proteins use dispersed NLS signals (to be published elsewhere). A very large effort was required to delineate the NLS signals from several *P. sojae* nuclear proteins to the point that a small highly efficient NLS tag could be constructed (to be published elsewhere). A second hurdle was that no RNA polymerase III promoters had been characterized in oomycetes, and that the U6 genes we attempted to use did not appear to be well transcribed under our transformation conditions. This we solved by using a strong RNA polymerase II promoter (*RPL41*) in conjunction with self-cleaving ribozymes to release the sgRNA from the RNA polymerase II transcript. A third hurdle was effective expression of the nuclease protein. Out initial attempts to use TALENs were blocked because the TALEN constructs were silenced extremely strongly in *P. sojae*, presumably due to their highly repetitive nature. The humanized SpCas9 protein was readily expressed, but unexpectedly the hSpCas9-GFP fusion protein we used to validate expression was not effective in generating mutations in *P. sojae*. despite the fact that Cas9-GFP fusions have been used in other organisms such as *Toxoplasma gondii* (Shen *et al.*, 2014). We were successful only after we used the non-fused hSpCas9. A fourth hurdle was identifying effective sgRNAs. One of the two sgRNAs predicted using the *sgRNA Designer* web tool proved ineffective both *in vitro* and *in vivo*. This sgRNA, sgRNA-A, was found to be strongly self-complementary, potentially preventing hybridization with the target DNA. This observation, which is in agreement with Peng *et al.*, (2015), underlines that sgRNAs with strong secondary structure predictions should be be eliminated. We also observed a discrepancy between *in vitro* and *in vivo* assays of Cas9/sgRNA activity, namely that the sgRNA lacking ribozymes was functional in the *in vitro* assay, whereas only the sgRNA flanked by ribozymes was effective *in vivo*. There clearly is room for further optimization of sgRNA design.

Using a single sgRNA, we were able to generate small indels at the site of the Cas9 cleavage. While these were useful, future attempts to disrupt genes with large deletions will likely require a pair of sgRNAs, or else the use of homology-directed repair to introduce specific mutations or to replace the gene entirely, as we did in replacing *Avr4/6* with *NPT II*. Of interest is the fact that from two of our transformant lines we were able to recover two or three different mutations. Several of these mutations were segregated into different single zoospore lines, indicating that the mutations had occurred in different nuclei of the regenerating protoplasts. One set of lines were biallelic indicating that two different mutations had occurred in the same nucleus. The fact that relatively few distinct mutations were recovered in each case might be explained if the mutations occurred very early during protoplast regeneration when there were limited numbers of nuclei present. Alternatively, the recovered mutations may have been the ones that occurred the earliest and therefore the nuclei carrying them came to dominate the heterokaryotic transformant. Also of interest is the observation that most of the mutations were recovered as homozygotes in this diploid organism. We speculate that once a mutation occurred in one allele that made it resistant to further Cas9 cleavage, cleavage of the remaining unmodified allele by Cas9 in most cases led to gene conversion of that allele to match the first allele.

It was reported that in some cases over-expression of Cas9 negatively impacts the growth of the organism, e.g. the parasite *Trypanosoma cruzi* (Peng *et al.*, 2015). However, during sub-culture and infection assays of all our Cas9-expressing mutants, we did not observe negative effects on *P. sojae* growth nor on overall virulence.

CRISPR-mediated gene disruptions and gene replacements will find numerous applications in *P. sojae* and other oomycetes. Gene disruptions will enable genes to be knocked out much more surely than by silencing. With careful design of sgRNAs, single members of closely related gene families could be eliminated, or else tags for transcriptional measurements could be introduced, to determine their individual contributions. Alternatively, entire gene clusters could be eliminated with a pair of sgRNAs. Gene replacements will enable mutations of all kinds, including promoter mutations and epitope and fluorescent protein tags, to be introduced into the endogenous gene where expression and phenotypes will not be confounded by position effects, over-expression artifacts, and contributions from the unmodified native gene. Targeted gene insertions are also expected to solve a longstanding problem in some oomycetes such as *P. sojae* where ectopic transgenes are invariably poorly expressed, even from the strongest promoter. Gene disruptions will also be useful for creating a much wider choice of selectable markers for transformation, through the creation of auxotrophic mutants. Disruption of the *Ku70* and/or *Ku80* genes required for non-homologous end joining may also result in higher efficiencies of homologous gene replacement (Arentshorst *et al.*, 2012). Gene deletions could also be used to remove integrated transgenes so that selectable markers such as *NPT II* can be recycled for repeated transformation experiments.

Some of the advantages of these CRISPR-enabled approaches are illustrated by insights gained from our manipulation of the *Avr4/6* gene. The elimination of the gene by replacement with *NPT II* confirms that *Avr4/6* makes a major contribution to recognition of the pathogen by plants containing the *Rps*4 and *Rps*6 loci, consistent with the findings of Dou *et al.*, (2010). However, the mutants did not kill the *Rps*4- and *Rps*6-containing plants as completely as they killed *rps* plants lacking *Rps*4 and *Rps*6. The *Rps*4 and *Rps*6-containing isolines were produced by introgression, not by transformation with individual *R* genes (Sandhu *et al.*, 2004). Since the *Rps*4 and *Rps*6 loci, which are allelic, both contain many NB-LRR genes (Sandhu *et al.*, 2004), we speculate that additional NB-LRR genes at these loci (or even the *Rps*4 and *Rps*6 genes themselves) can recognize additional effectors produced by the *P. sojae* strain used in these studies (P6497). Dou *et al* (2010) also observed that *P. sojae* strains silenced for Avr4/6 did not completely kill *Rps*4 and *Rps*6-containing lines, but those observations were ascribed to incomplete silencing of *Avr4/6*. With the availability of complete *Avr4/6* deletion mutants, we can now more confidently conclude that the *Rps*4 and *Rps*6 loci make additional contributions to resistance other than through recognition of Avr4/6.

The two frameshift mutants killed *Rps*6 plants nearly as well as the HDR mutants (62% combined killing versus 74%) indicating that these mutants were no longer recognized by *Rps*6-containing plants. T32-3 killed *Rps*4 plants better than the two HDR mutants (83% versus 74% killing), but T11-2-5 was clearly avirulent on the *Rps*4 plants (40% versus 74%; WT = 31%). The explanation for this difference may lie in the observation by Dou *et al* (2010) that the N-terminal domain of Avr4/6, up to and including the dEER motif, is sufficient for recognition by *Rps*4 plants, whereas recognition by *Rps*6 plants requires the C-terminus. Since the site of the CRISPR-induced NHEJ mutations is immediately upstream of the dEER motif, the +1 frameshift in T32-3 retained the N-terminal domain but eliminated the dEER motif, presumably abolishing effector entry into the plant (Dou *et al.*, 2008b). The -1 frameshift in T11-2-5 also eliminated the dEER motif, but in its place created a highly positively charged sequence (RKKRNARSR). Thus we speculate that this positively charged sequence may act as a surrogate cell entry sequence (Snyder & Dowdy, 2004, Dou *et al.*, 2008b, Kale *et al.*, 2010, Milletti, 2012) delivering the N-terminal fragment for recognition by Rps4.

Of the three in-frame deletions, the two single amino acid deletion mutants scored as intermediate to avirulent on both *Rps*4 and *Rps*6 plants, suggesting that recognition was only slightly impaired. The two amino acid deletion mutant (T11-1) however was actually slightly more virulent on *Rps*4 plants than the HDR mutants (86% versus 74% killing) while slightly less virulent on the *Rps*6 plants (64%). We speculate that the two residue deletion may have disrupted the structure of Avr4/6 with possibly a stronger effect on the immediate region of the N-terminus required for recognition by *Rps*4 plants.

In summary, the adaptation of CRISPR/Cas9-mediated gene targeting to oomycetes is expected to rapidly advance the functional analysis of these extremely destructive and important plant and animal pathogens.

## EXPERIMENTAL PROCEDURES

### *Phytophthora sojae* strains and growth conditions

The reference *P. sojae* isolate P6497 (Race 2) used in this study was routinely grown and maintained in cleared V8 medium at 25 °C in the dark. Zoospores were induced and isolated from ∼ 1-week old cultures grown on clarified V8 agar, as previously described (Judelson *et al.*, 1993a). Single *P. sojae* transformants were incubated in 12-well plates containing V8 media supplemented with 50 µg/mL G418 (Geneticin) for 2∼3 days before genomic DNA was isolated by miniprep.

### sgRNA design

sgRNA target sites were selected according to the web-tool *sgRNA Designer* (http://www.broadinstitute.org/rnai/public/analysis-tools/sgrna-design; (Doench *et al.*, 2014). Potential off-target sites were checked using the FungiDB (www.fungidb.org) alignment search tool (BLASTN) against the *P. sojae* genome and visual inspection of the results. Sequences that perfectly matched the final 12 nt of the target sequence and NGG PAM sequence were discarded (Cong *et al.*, 2013). Ribozymes were designed according to Gao, *et al.* (2014). The first six nucleotides of the Hammerhead (HH) ribozyme were designed to be the reverse complement of the first six nucleotides of the sgRNA target sequences.

### Plasmid construction

All the primers used in this study are listed in the Table S1 in the supplemental material. A map and sequence file for the plasmid backbones used for expressing Cas9 and the sgRNAs targeting the *Avr4/6* locus can be found in Fig. S4 in the supplemental material. To express *hSpCas9* and sgRNAs effectively in *P. sojae*, we first created a new *Phytophthora* expression plasmid backbone pYF2 by combining elements from pHamT34 (Judelson *et al.*, 1991), pUN (Dou *et al.*, 2008a) and pGFPN (Ah-Fong & Judelson, 2011) as follows. (i) The *HSP70* terminator was PCR amplified from pGFPN by using primers BlHSP70T_AgeI_F and BlHSP70T_NsiI_EcoRI_R which added an *Eco* RI site, and was then inserted into pUN using *Kpn* I and *Nsi* I sites, placing the *NPT II* gene under the control of the *RPL41* promoter and *HSP70* terminator. (ii) The entire *NPT II* cassette from (i) was then extracted by digestion with *Eco* RI and inserted into the *Eco* RI site of pHAMT34, resulting in pYF1. (iii) A synthetic multiple cloning site fragment containing the restriction enzyme sites *Xma I-Cla* I-*Stu* I-*Sac* II-*Spe* I-*Bsi* WI-*Afl* II-*Kpn* I was introduced between the *Xma* I and *Kpn* I sites of pYF1 by oligo annealing, creating pYF2. The plasmid pYF2-GFP was generated by adding an *eGFP* fragment amplified from pGFPN using primers GFP_AflII_F and GFP_ApaI_R into the restriction enzyme sites *Afl* II and *Apa* I of pYF2. The PsNLS, reported in Fang & Tyler (2015), was tested using the plasmid pYF2-PsNLS-2XGFP, which was constructed by two steps. (i) An extra *GFP* was amplified from the same plasmid pGFPN using primers GFP_SpeI_F and GFP_Afl II_R and inserted into the restriction sites *Spe* I and *Afl* II of the plasmid pYF2-GFP, generating pYF2-2XGFP. (ii) The PsNLS was inserted by annealing of two oligonucleotides encoding the NLS (MHKRKREDDTKVRRRMHKRKREDDTKVRRRMHKRKREDDTKVRRR).

To generate *hSpCas9-*expression plasmids pYF2-PsNLS-*hSpCas9*-GFP and pYF2-PsNLS-*hSpCas9*, the coding fragment of *hSpCas9* nuclease from the plasmid pSpCas9 (BB)-2A-GFP (PX458, Addgene plasmid # 48138) were subcloned into the *Spe* I and *Afl* II sites or *Spe* I and *Apa* I sites (there is an *Apa* I site in *hSpCas9*, so the two digested fragments were cloned sequentially), respectively, of plasmid pYF2-PsNLS-2XGFP containing the *P. sojae* NLS.

To generate plasmids pYF2.2-GFP-sgRNA and pYF2.2-GFP-Ribo-sgRNA, the plasmid pYF2-GFP was first modified to pYF2.2-GFP by mutating the unexpected *Age* I site in the junction of *eGFP* and the Ham34 terminator to a *Bsp* EI site through QuikChange Lightning Multi Site-Directed Mutagenesis (Agilent Technologies), and the *Kpn* I site was also removed. Then sgRNAs with or without ribozyme were PCR amplified from the same plasmid, PX458, and inserted into the sites *Age* I and *Nhe* I of pYF2.2-GFP.

To generate the construct for replacing the entire ORF of *Avr4/6*, the *NPT II* coding region together with 250 bp, 500 bp or 1 kb of 5′ and 3′ flanking regions outside the *Avr4/6* coding region were PCR amplified and cloned into the plasmid pBluescript II KS+ by In-Fusion^®^ HD Cloning Kit (Clontech).

### sgRNA:Cas9 *in vitro* activity assay

To test the activity of the designed sgRNAs, an *in vitro* cleavage assay was carried out as previously described (Gao & Zhao, 2014). Briefly, sgRNA was *in vitro* transcribed through run-off reactions with T7 RNA polymerase using the MEGAshortscript^TM^ T7 kit (Ambion) according to the manufacturer’s manual. Templates for sgRNA synthesis were generated by PCR amplification from the sgRNA expression plasmid pYF2.2-sgRNAs (Table S1). The target DNA was amplified from pCR2.1-Avr4/6 using primer M13F and M13R (Supplemental material). SpCas9 nuclease was purchased from New England Biolabs, and the cleavage assay was performed according to the product manual.

### Improved transformation of *P. sojae*

Polyethylene glycol (PEG) mediated protoplast transformations were conducted to introduce DNA into *P. sojae*. To achieve a higher transformation rate, we improved the previously described transformation methods (Mcleod *et al.*, 2008, Dou *et al.*, 2008a) as follows. 2-4 days old *P*. *sojae* mycelial mats, cultured in nutrient pea broth, were harvested and pre-treated with 0.8 M mannitol for 10 min, then digested in 20 mL enzyme solution (0.4 M mannitol, 20 mM KCl, 20mM MES, pH 5.7, 10 mM CaCl_2_, 0.5% Lysing Enzymes from *Trichoderma harzianum* [Sigma L1412], and 0.5% CELLULYSIN^®^ Cellulase [Calbiochem 219466]) for ∼40 min at room temperature with gentle shaking. The mixture was filtered through a Falcon™ Nylon Mesh Cell Strainer (BD Biosciences) and protoplasts were pelleted by centrifugation at 1,200g for 2 min in a Beckman Coulter benchtop centrifuge with swing buckets. After washing with 30 mL W5 solution (5 mM KCl, 125 mM CaCl_2_, 154 mM NaCl, and 177 mM glucose), protoplasts were resuspended in 10 mL W5 solution and left on ice for 30 min. Protoplasts were collected by centrifugation at 1,200 g for 2 min in the Beckman centrifuge and resuspended at 10 /mL in MMg solution (0.4 M mannitol, 15 mM MgCl_2_ and 4 mM MES, pH 5.7). DNA transformation was conducted in a 50 mL Falcon tube, where 1 mL protoplasts were well mixed with 20-30 µg DNA for single plasmid transformation. For co-transformation experiments, 20-30 µg of the plasmid carrying the *NPT II* selectable marker gene was used, together with an equimolar ratio of any other DNAs included. Then, three successive aliquots of 580 µl each of freshly made polyethylene glycol (PEG) solution (40% PEG 4000 v/v, 0.2 M mannitol and 0.1 M CaCl_2_) were slowly pipetted into the protoplast suspension and gently mixed. After 20 min incubation on ice, 10 mL pea broth containing 0.5 M mannitol were added, and the protoplasts were regenerated overnight at 18°C in the dark. For production of stable transformants, the regenerated protoplasts were collected by centrifugation at 2,000 g for 2 min in the Beckman centrifuge, and then resuspended and evenly divided into three Falcon tubes containing 50 mL liquid pea broth containing 1% agar (42 ºC), 0.5 M mannitol and 50 µg/mL G418 (AG Scientific). The resuspended protoplasts were then poured into empty 60 mm × 15 mm petri dishes. Mycelial colonies could be observed after 2 d incubation at 25°C in the dark. The visible transformants were transferred to V8 liquid media containing 50 µg/mL G418 and propagated for 2∼3d at 25 ºC prior to analysis. For transient expression, 50 µg/mL G418 was usually added into the regeneration medium after overnight recovery, to enrich the positive transformants. After 1 d incubation at 25 ºC in the dark, hyphae were collected for genomic DNA extraction.

### Detection and quantification of targeted mutagenesis

To detect the results of targeted mutagenesis in transformants, total genomic DNA (gDNA) was extracted from pooled or individual *P. sojae* transformants. For pooled transformants, 48 h after transformation 1 mL of the mycelial culture was pelleted, resuspended in 500 mL lysis buffer (200 mM Tris, pH 8.0, 200 mM NaCl, 25 mM EDTA, pH 8.0, 2% SDS, plus 0.1 mg/mL RNase added prior to use) and broken by vortexing with 0.5 mm glass beads. For individual transformants, approximately a 7 mm diameter clump of *P. sojae* hyphae were blotted dry on Kimwipe™ paper, then frozen in liquid nitrogen. Frozen mycelia was transferred into a 1.5 mL Eppendorf tube and ground to a powder using a polypropylene pestle, then resuspended in 500 µL lysis buffer. Hyphal lysates were incubated at 37 ºC, 30 min for RNA digestion. gDNA mixture was extracted with an equal volume of phenol- chloroform- isoamyl alcohol (25:24:1, pH 8.3), and then with an equal volume of chloroform. The aqueous phase was recovered and precipitated by adding 0.5 volume of isopropanol. After being pelleted and washed with 70% ethanol, the gDNA was dried under vacuum and dissolved in 40 µL water.

All PCR amplifications were conducted using Phusion® high-fidelity DNA polymerase (New England Biolabs) in order to exclude the possibility of mutations causing during PCR amplification. Generally ∼10 ng gDNA was used as PCR template. Nested PCR was conducted if necessary, using 1:1000 diluted PCR products as a DNA template for the second round.

To detect NHEJ mutations, the entire 372 bp *Avr4/6* ORF was amplified and examined by digestion with the relevant restriction enzyme. For pooled transformants, a nested PCR was performed to enrich the mutated target before sequencing; this step was not needed for individual transformants, including single zoospore lines. To detect HDR events in pooled and individual transformants (other than zoospore lines), primers located outside the *Avr4/6* homology arms and in *NPT II* gene were used. For screening single zoospore lines, PCR was performed by only using primers outside the homology arms. In both cases, nested PCR was carried out for efficient amplification of the targets. PCR products were sequenced directly by the Sanger dideoxy method in the Oregon State University Center for Genome Research and Biocomputing.

### Confocal Microscopy

Laser scanning confocal microscopy (Zeiss LSM 780 NLO) was used to examine the expression and subcellular localization of hSPCas9 fused to the NLS and to GFP. Living hyphae were picked from liquid cultures after 2-3 days growth and regeneration of transformants. Samples were stained with DAPI (4’, 6-diamidino-2-phenylindole) for 20 min in the dark (Talbot, 2001) before microscopy examination. Images were captured using a 63X oil objective with excitation/emission settings (in nm) 405/410-490 for DAPI, and 488/510∼535 for GFP.

### Infection assays

The ability of *Avr4/6* mutants to infect soybean plants carrying *Rps4* and *Rps6* was evaluated by hypocotyl inoculation as previously described (Dou *et al.*, 2010). The wild type and mutant *P.sojae* strains were grown on V8 plates without G418 selection for ∼5 days. Seeds of soybean cultivars HARO(1-7)1 *(rps)*, HARO4272 (Harosoy background, *Rps4, Rps7)*, HARO6272 (Harosoy background, *Rps6, Rps7)*, Williams *(rps)*, L85-2352 (Williams background, *Rps4)*, and L89-1581 (Williams background, *Rps6)* were provided by M. A. Saghai-Maroof (Virginia Tech). Each pathogenicity test was performed in triplicate, each replicate consisting of at least 19 seedlings. A strain was considered avirulent if the number of inoculated *Rps4* or *Rps6* seedlings surviving was significantly higher than among the seedlings lacking the *Rps* gene, as determined by Fisher’s exact test (Sokal & Rohlf, 1995) and the number was not significantly different than the number of surviving seedlings inoculated with the unmodified control strain P6497. A strain was considered virulent if the number of inoculated *Rps4* or *Rps6* seedlings surviving was not significantly different than among the seedlings lacking the *Rps* gene, and the number was significantly fewer than the number of surviving seedlings inoculated with the unmodified control strain P6497. A strain was considered intermediate if the number of inoculated *Rps4* or *Rps6* seedlings surviving was not significantly different than among the seedlings lacking the *Rps* gene, and the number was also not significantly different than the number of surviving seedlings inoculated with the unmodified control strain P6497. A strain was also considered intermediate if the number of inoculated *Rps4* or *Rps6* seedlings surviving was significantly higher than among the seedlings lacking the *Rps* gene, and the number was also significantly lower than the number of surviving seedlings inoculated with the unmodified control strain P6497. Intermediate phenotypes were further designated intermediate/virulent or intermediate/avirulent if the p values indicating a difference from virulent or avirulent controls differed by more than 10 fold.

## ACKNOWLEDGEMENTS

We thank F. Arredondo, S. Taylor and D. Wellappili (Oregon State University) for experiment assistance, M.A. Saghai-Maroof (Virginia Tech) for soybean seed, and H. Judelson (UC-Riverside) and members of the Tyler Laboratory for useful advice. We acknowledge the Sequencing and Confocal Microscopy Facilities of the Center for Genome Research and Biocomputing at Oregon State University. This work was supported in part by grant 2011-68004-30104 from the Agriculture and Food Research Initiative of the USDA National Institute for Food and Agriculture.

## SUPPLEMENTARY INFORMATION

Supplementary Table 1 The sequences of oligonucleotides used in this study

Supplementary Fig. S1 *Phytophthora* U6 promoter evaluation

Supplementary Fig. S2 *Phytophthora* U6 promoter evaluation

Supplementary Fig. S3 Representative sequencing chromatograms of the Avr4/6 mutations in the single zoospore-purified mutants

Supplementary Fig. S4 Plasmid backbones used for expression of *hSpCas9* and sgRNA in *P. sojae*

Supplementary Methods

Supplementary Sequence Information

**Supplemental Table 1.**
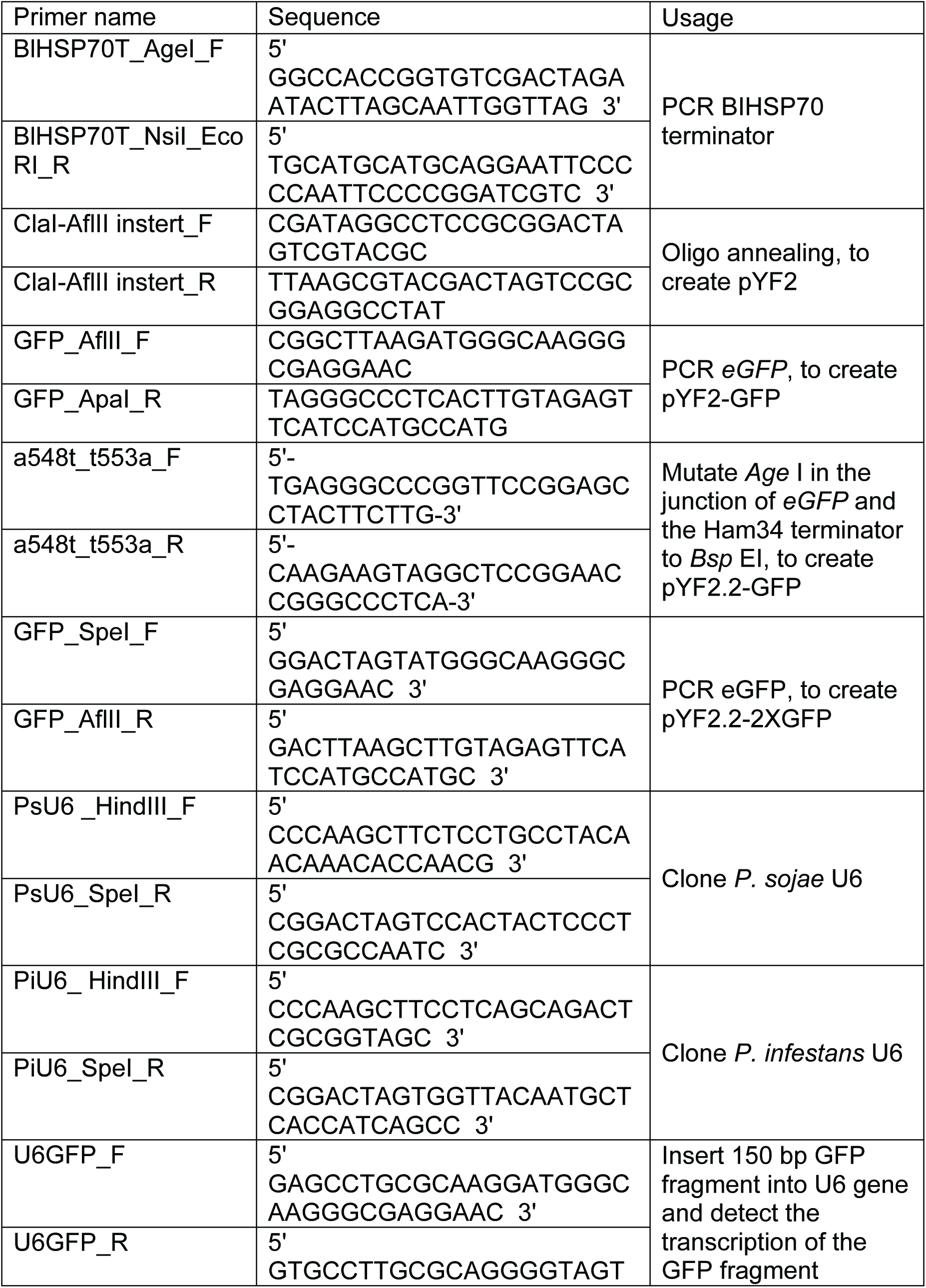
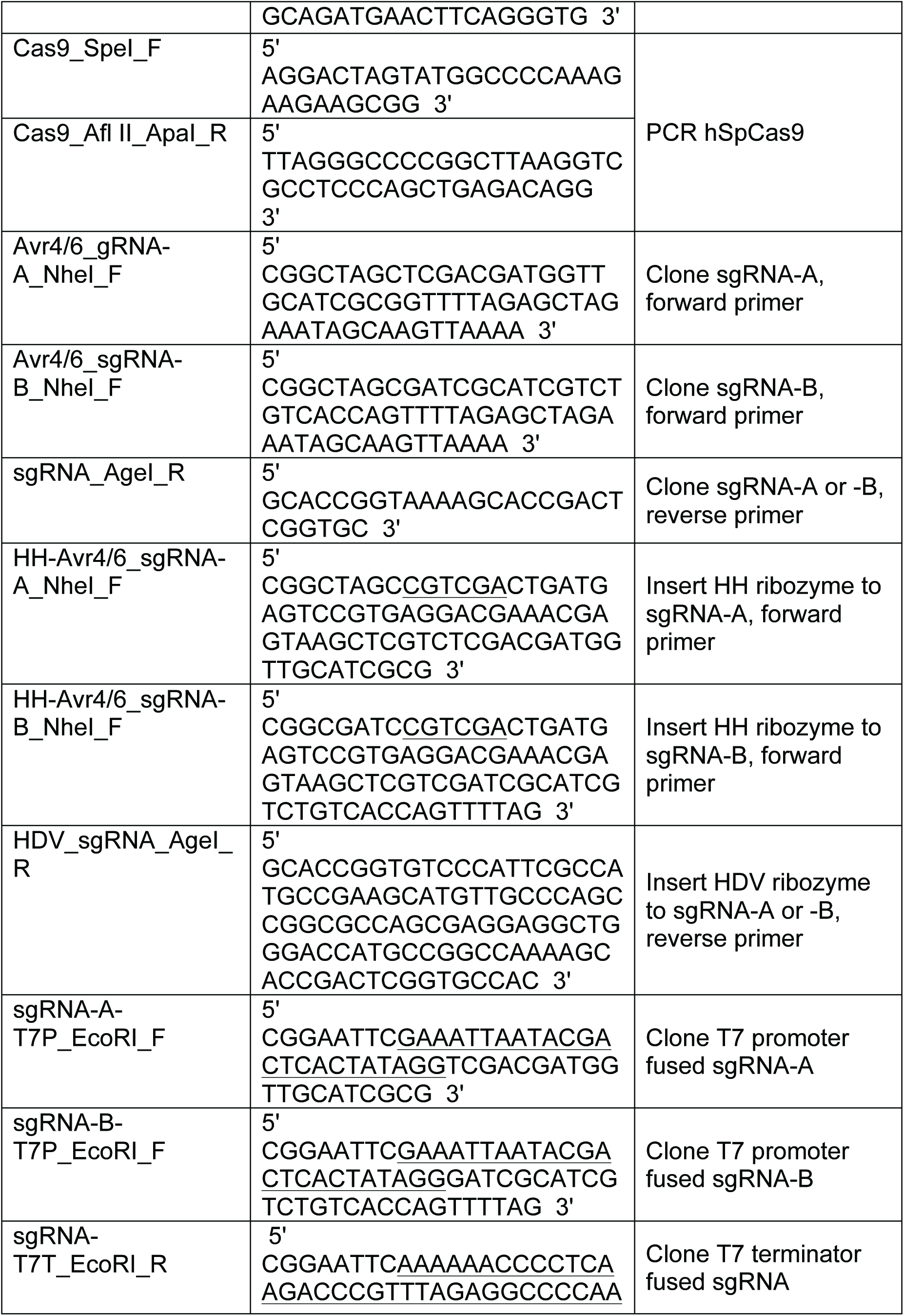
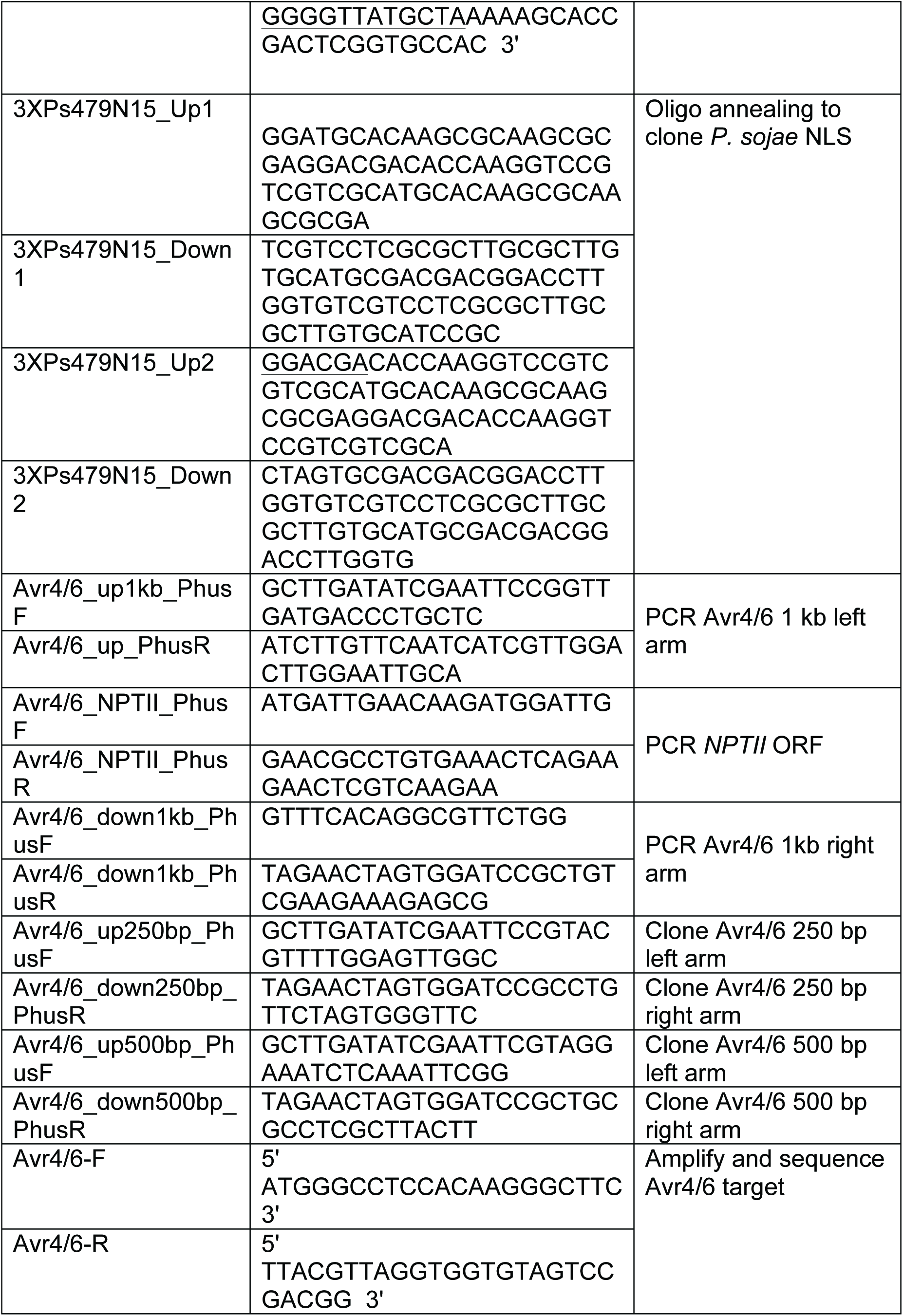
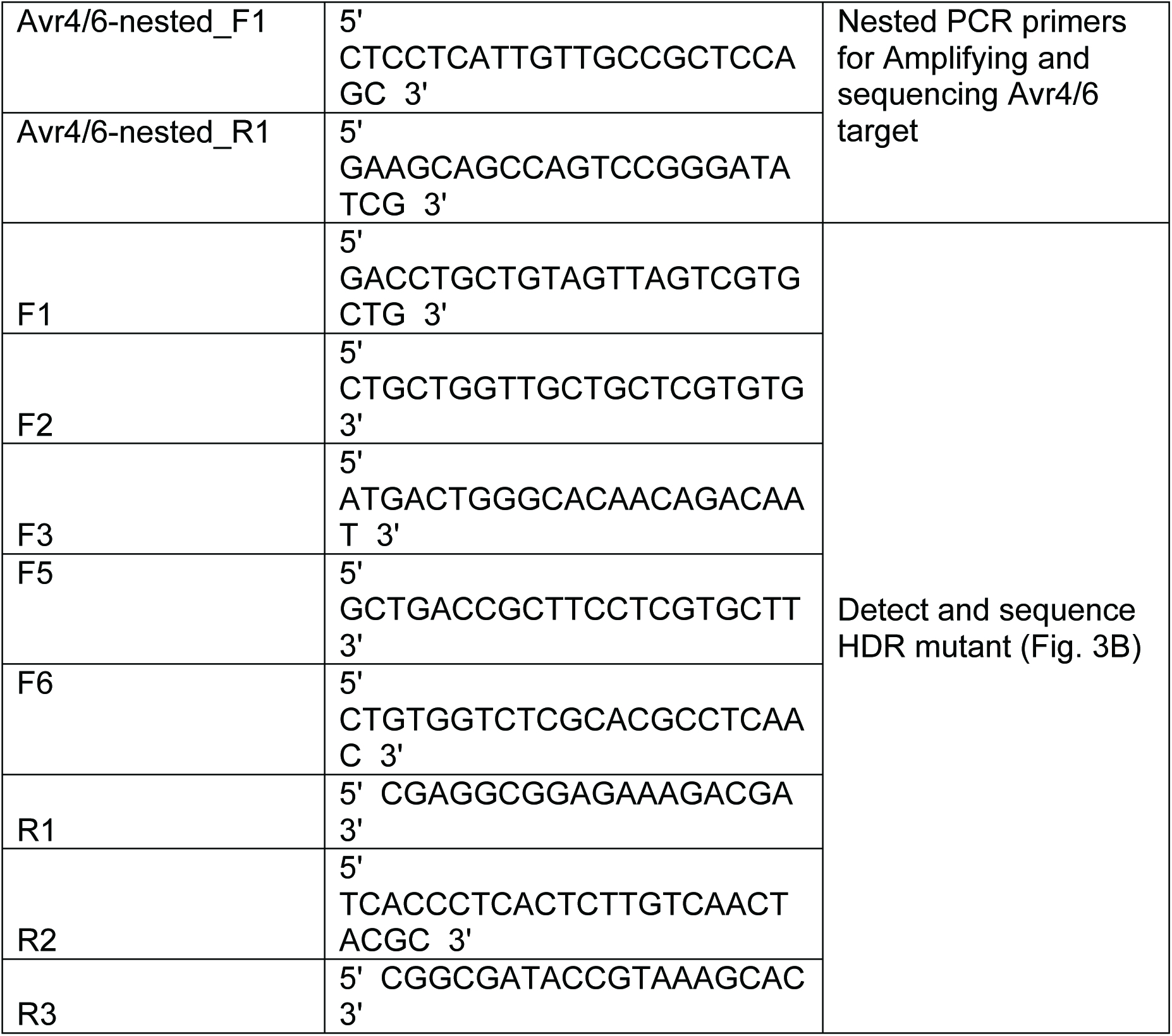
The sequences of oligonucleotides used in this study

**Fig. S1.**
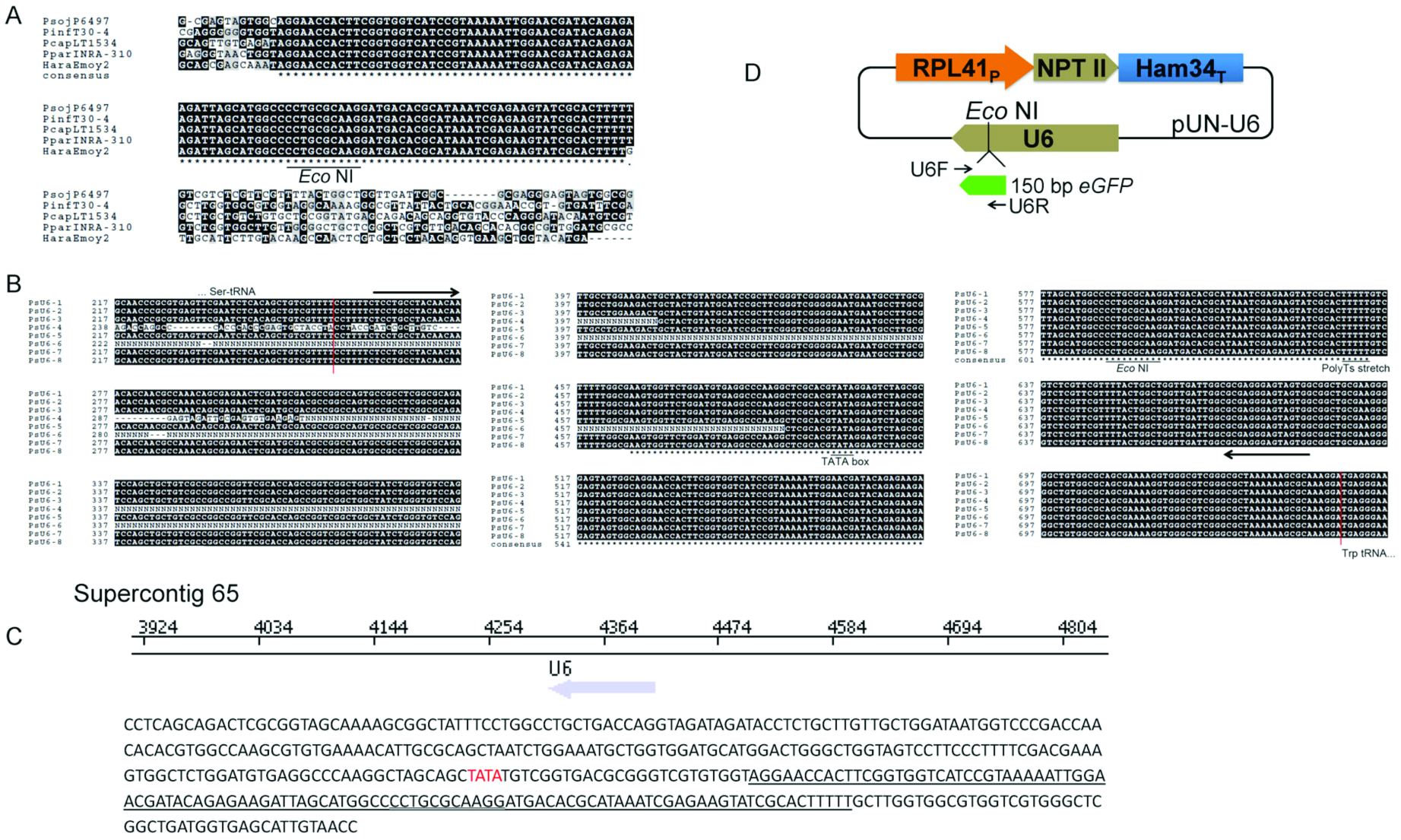
*Phytophthora* U6 promoter evaluation. (A) Alignment of selected oomycete U6 genes, showing that U6 transcripts are highly conserved. The numbers of U6 genes were variable in different oomycete species, 8 in *P. sojae P6497* (*PsojP6497*), 127 in *P. infestans* (*PinfT30-4*), 5 in *capsici LT1534* (*PcapLT1534*), 1 *in P.* parasitica (Ppar INRA-310), and 1 in *H. arabidopsidis (HaraEmoy2)* respectively. Genome data are obtained from the fungidb.org website. (B) Alignment of the 8 annotated *P. sojae* U6 genes. PsU6-1 was used to test promoter activity. The red lines indicate the border of the upstream and downstream tRNAs repectively. (C) One of the 127 *P. infestans* U6 genes cloned to test U6 promoter activity. (Top) Position of the PiU6 gene on *Phytophthora infestans* T30-4 Supercontig 65. (Bottom) PiU6 sequence used for promoter activity test. The putative U6 coding region is underlined; a putative TATA-box is in red. (D) The plasmid used for testing the functions of the PsU6-1 and PiU6 promoters in *P. sojae*. Residues 1-150 bp of *eGFP* was used as a transcription detection maker. Arrows indicate the primer pair U6GFP_F and U6GFP_R used for cloning the *eGFP* fragment and also for detection of U6 transcripts by RT-PCR. The *Eco* NI restriction enzyme site used for inserting the GFP detection marker is underlined in (A) and (B) and double underlined in (C).

**Fig. S2.**
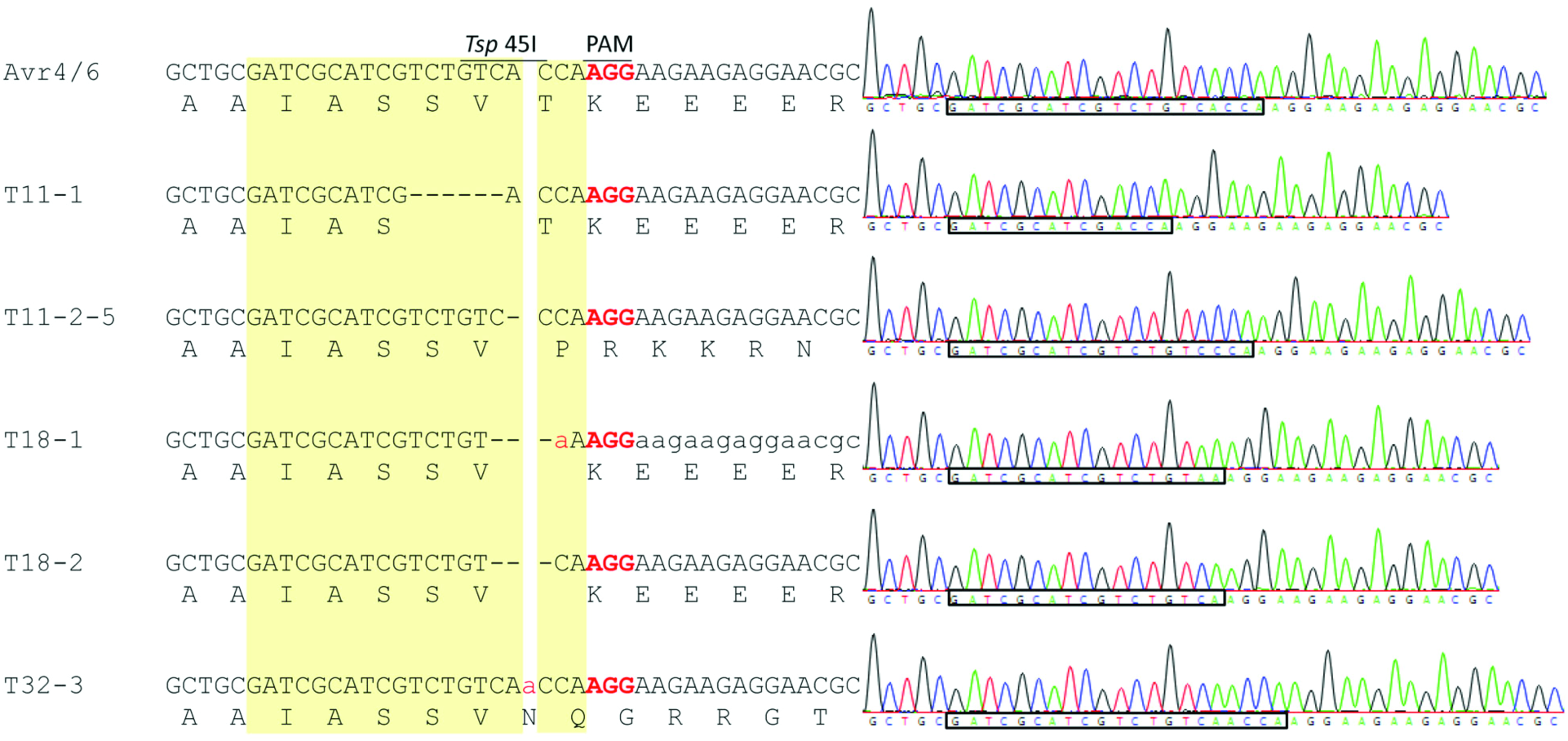
Representative sequencing chromatograms of the Avr4/6 mutations in the single zoospore-purified mutants. Regions in box showing the sgRNA target sites within the *Avr4/6* gene. The unambiguous sequencing profiles indicate that these mutant lines are all homozygous.

**Fig. S3.**
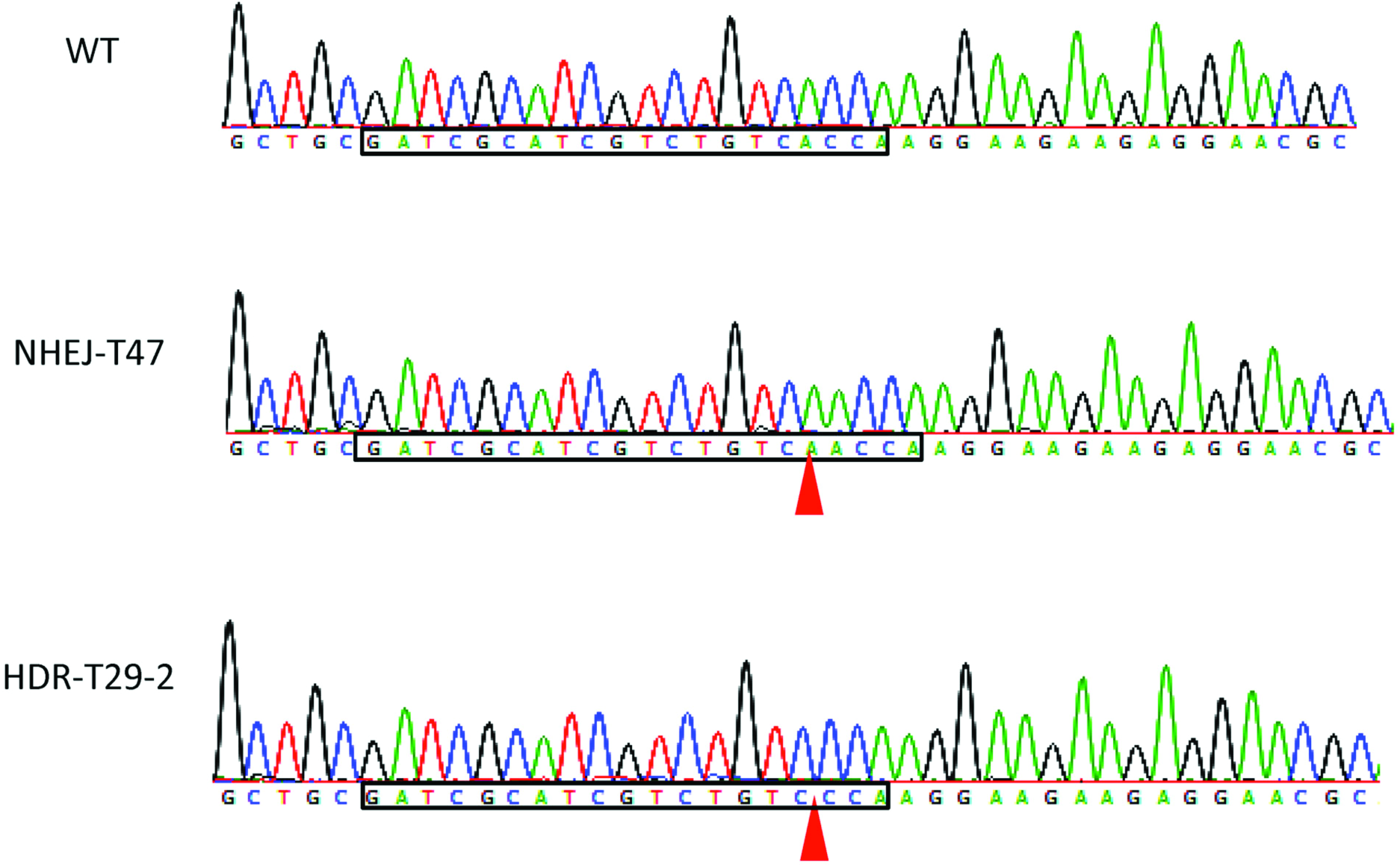
Sanger sequencing profiles revealing that the sub-cultured Cas9:sgRNA transformant T47 (NHEJ-T47) and HDR mutant T29 (HDR-T29-2) had NHEJ mutations (one bp insertion and deletion respectively). Red triangles indicate the differences between wild type (WT) and mutants.

**Fig. S4.**
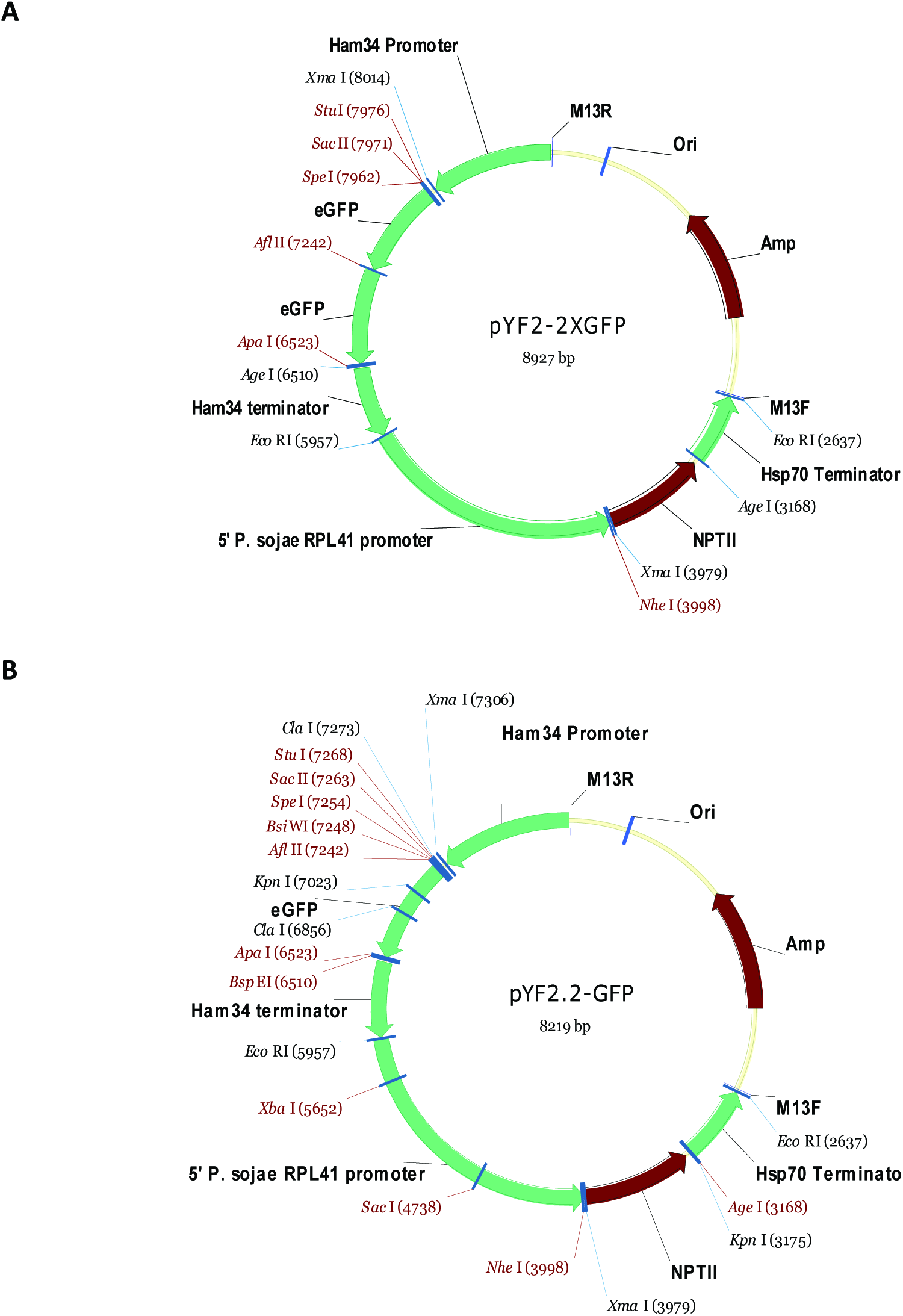
Plasmid backbones used for expression of *hSpCas9* and sgRNA in *P. sojae*. (A) pYF2-2XGFP is used for tracking the subcellular localization and expression of PsNLS fused *hSpCas9*. PsNLS is inserted into *Sac* II and *Spe* I sites. *hSpCas9* is inserted into *Spe* I and *Afl* II sties for subcellular localization examination and *Spe* I and *Apa* I sites for CRISPR expression. (B) pYF2.2-GFP is used for expression of sgRNA (inserted into *Nhe* I and *Age* I sites).

**>pYF2.2-GFP**

AAGCTTGGCGTAATCATGGTCATAGCTGTTTCCTGTGTGAAATTGTTATCCGCTCACAATTCCACACAACATACGAGCCGGAAGCATAAAGTGTAAAGCCTGGGGTGCCTAATGAGTGAGCTAACTCACATTAATTGCGTTGCGCTCACTGCCCGCTTTCCAGTCGGGAAACCTGTCGTGCCAGCTGCATTAATGAATCGGCCAACGCGCGGGGAGAGGCGGTTTGCGTATTGGGCGCTCTTCCGCTTCCTCGCTCACTGACTCGCTGCGCTCGGTCGTTCGGCTGCGGCGAGCGGTATCAGCTCACTCAAAGGCGGTAATACGGTTATCCACAGAATCAGGGGATAACGCAGGAAAGAACATGTGAGCAAAAGGCCAGCAAAAGGCCAGGAACCGTAAAAAGGCCGCGTTGCTGGCGTTTTTCCATAGGCTCCGCCCCCCTGACGAGCATCACAAAAATCGACGCTCAAGTCAGAGGTGGCGAAACCCGACAGGACTATAAAGATACCAGGCGTTTCCCCCTGGAAGCTCCCTCGTGCGCTCTCCTGTTCCGACCCTGCCGCTTACCGGATACCTGTCCGCCTTTCTCCCTTCGGGAAGCGTGGCGCTTTCTCAATGCTCACGCTGTAGGTATCTCAGTTCGGTGTAGGTCGTTCGCTCCAAGCTGGGCTGTGTGCACGAACCCCCCGTTCAGCCCGACCGCTGCGCCTTATCCGGTAACTATCGTCTTGAGTCCAACCCGGTAAGACACGACTTATCGCCACTGGCAGCAGCCACTGGTAACAGGATTAGCAGAGCGAGGTATGTAGGCGGTGCTACAGAGTTCTTGAAGTGGTGGCCTAACTACGGCTACACTAGAAGGACAGTATTTGGTATCTGCGCTCTGCTGAAGCCAGTTACCTTCGGAAAAAGAGTTGGTAGCTCTTGATCCGGCAAACAAACCACCGCTGGTAGCGGTGGTTTTTTTGTTTGCAAGCAGCAGATTACGCGCAGAAAAAAAGGATCTCAAGAAGATCCTTTGATCTTTTCTACGGGGTCTGACGCTCAGTGGAACGAAAACTCACGTTAAGGGATTTTGGTCATGAGATTATCAAAAAGGATCTTCACCTAGATCCTTTTAAATTAAAAATGAAGTTTTAAATCAATCTAAAGTATATATGAGTAAACTTGGTCTGACAGTTACCAATGCTTAATCAGTGAGGCACCTATCTCAGCGATCTGTCTATTTCGTTCATCCATAGTTGCCTGACTCCCCGTCGTGTAGATAACTACGATACGGGAGGGCTTACCATCTGGCCCCAGTGCTGCAATGATACCGCGAGACCCACGCTCACCGGCTCCAGATTTATCAGCAATAAACCAGCCAGCCGGAAGGGCCGAGCGCAGAAGTGGTCCTGCAACTTTATCCGCCTCCATCCAGTCTATTAATTGTTGCCGGGAAGCTAGAGTAAGTAGTTCGCCAGTTAATAGTTTGCGCAACGTTGTTGCCATTGCTACAGGCATCGTGGTGTCACGCTCGTCGTTTGGTATGGCTTCATTCAGCTCCGGTTCCCAACGATCAAGGCGAGTTACATGATCCCCCATGTTGTGCAAAAAAGCGGTTAGCTCCTTCGGTCCTCCGATCGTTGTCAGAAGTAAGTTGGCCGCAGTGTTATCACTCATGGTTATGGCAGCACTGCATAATTCTCTTACTGTCATGCCATCCGTAAGATGCTTTTCTGTGACTGGTGAGTACTCAACCAAGTCATTCTGAGAATAGTGTATGCGGCGACCGAGTTGCTCTTGCCCGGCGTCAATACGGGATAATACCGCGCCACATAGCAGAACTTTAAAAGTGCTCATCATTGGAAAACGTTCTTCGGGGCGAAAACTCTCAAGGATCTTACCGCTGTTGAGATCCAGTTCGATGTAACCCACTCGTGCACCCAACTGATCTTCAGCATCTTTTACTTTCACCAGCGTTTCTGGGTGAGCAAAAACAGGAAGGCAAAATGCCGCAAAAAAGGGAATAAGGGCGACACGGAAATGTTGAATACTCATACTCTTCCTTTTTCAATATTATTGAAGCATTTATCAGGGTTATTGTCTCATGAGCGGATACATATTTGAATGTATTTAGAAAAATAAACAAATAGGGGTTCCGCGCACATTTCCCCGAAAAGTGCCACCTGACGTCTAAGAAACCATTATTATCATGACATTAACCTATAAAAATAGGCGTATCACGAGGCCCTTTCGTCTCGCGCGTTTCGGTGATGACGGTGAAAACCTCTGACACATGCAGCTCCCGGAGACGGTCACAGCTTGTCTGTAAGCGGATGCCGGGAGCAGACAAGCCCGTCAGGGCGCGTCAGCGGGTGTTGGCGGGTGTCGGGGCTGGCTTAACTATGCGGCATCAGAGCAGATTGTACTGAGAGTGCACCATATGCGGTGTGAAATACCGCACAGATGCGTAAGGAGAAAATACCGCATCAGGCGCCATTCGCCATTCAGGCTGCGCAACTGTTGGGAAGGGCGATCGGTGCGGGCCTCTTCGCTATTACGCCAGCTGGCGAAAGGGGGATGTGCTGCAAGGCGATTAAGTTGGGTAACGCCAGGGTTTTCCCAGTCACGACGTTGTAAAACGACGGCCAGTGAATTCCCCCAATTCCCCGGATCGTCTCATAGCTGCTCAAGACGTTTAGCAGATTGAGGATGCAGATCTTCTACTAATGAAACTACTGCAAACTACCAACAAAGAGGCGACCACAAAATGGACAGCCTATGCCGAGATTTGTTTAAGATACTCGATTTGCCGTCCAATTTTCATCAAGACGAAACACAGTGCATCGAACTGTGTTGTAAAAATTCATTTATTCTTGCAATTGTGACTCTCAAAGATTGATTAAACTTGAAAGCTGGTGAAAGCTTTCCTTCTTTTGGCATGTTTGTTACTAAGGCGGCGTAAGCACAATAGGCCCAGACTCATGCAAACCAGCCATAGTGACCTTCTCACTATGTTTCGCTTGAGATTTACTAGACTCTCGGGCATCTTCACAAAAGATGACAACCGAATCAACAAATTAAGTCAACGCTAAATCAAATATTTAGAAAATAAATTTTATTGTCGAAAAAACTCTGCTTTGACAATTTTCAAACTAACCAATTGCTAAGTATTCTAGTCGACACCGGTACCTATCAGAAGAACTCGTCAAGAAGGCGATAGAAGGCGATGCGCTGCGAATCGGGAGCGGCGATACCGTAAAGCACGAGGAAGCGGTCAGCCCATTCGCCGCCAAGCTCTTCAGCAATATCACGGGTAGCCAACGCTATGTCCTGATAGCGGTCCGCCACACCCAGCCGGCCACAGTCGATGAATCCAGAAAAGCGGCCATTTTCCACCATGATATTCGGCAAGCAGGCATCGCCATGGGTCACGACGAGATCCTCGCCGTCGGGCATGCGCGCCTTGAGCCTGGCGAACAGTTCGGCTGGCGCGAGCCCCTGATGCTCTTCGTCCAGATCATCCTGATCGACAAGACCGGCTTCCATCCGAGTACGTGCTCGCTCGATGCGATGTTTCGCTTGGTGGTCGAATGGGCAGGTAGCCGGATCAAGCGTATGCAGCCGCCGCATTGCATCAGCCATGATGGATACTTTCTCGGCAGGAGCAAGGTGAGATGACAGGAGATCCTGCCCCGGCACTTCGCCCAATAGCAGCCAGTCCCTTCCCGCTTCAGTGACAACGTCGAGCACAGCTGCGCAAGGAACGCCCGTCGTGGCCAGCCACGATAGCCGCGCTGCCTCGTCCTGCAGTTCATTCAGGGCACCGGACAGGTCGGTCTTGACAAAAAGAACCGGGCGCCCCTGCGCTGACAGCCGGAACACGGCGGCATCAGAGCAGCCGATTGTCTGTTGTGCCCAGTCATAGCCGAATAGCCTCTCCACCCAAGCGGCCGGAGAACCTGCGTGCAATCCATCTTGTTCAATCATTGTTGCCCGGGTGGATGCTCAGATGCTAGCGTCACCGAACGAATTAAAACTATAGTAAGTAAAGACTGTCGATGTAGCCATGTGGGGCCAATAGCTGCGAGGCTCAGAGAGAGACAAGGGGGTCGCCACTTACCCCGAACGATGTTCCAAGCAGAGCAGTCAGCAGAAAGTGAGGCTTGTTGCTGTCGGCGAGCGCAGCCTTGGGATCAACATGCGAGAGGATAGTTGGCTGCGAGCTTGCCAGCGTGGAGAGACTTGGTTATTGTAGGCAACAAGTCCCGCATGAAGGGCGTTGTATGGTGCAAAAAAGAGGAGATGGAATCTTCAAACACTTTGATTGGCTGGACAGCGGGCGCGGACTACTGGCCAACCAAAATGTGACCCCACTTGAGTGACACTTTCCCCACTATAGTGTCTTCAACTTGCACTGCAAGTGAAGTTGTTCTTGTTATTGCACTCGTTCTTACGTTATGGCTAGTACGCGCAGATGCAGAGCCTCCTCGGTGGTATTGCACGCCAACGAACCTCCATTCTCTGCGACATAAGGCTGTGGCAGTGCGCCCAGGAACTCGTAGAGAATGGTCAACTGCGCCGTCACCACTCCCGTGTTCGAGTCTTCCGTCGTGACGAGGGCCGACTTCACCGTCGCCACATTCGACAGCGCAGTGCGGTACATCTCCATCAGCATCGTGTCCAGTTGGGCTGTCACTTCCGTCGCGTTCGCCGTGAAGTCCTCGACAAAACTGTACGCCCAGTTGCCGAGCTCGAAGCTGACGCTGGCGAATCCGCTCGCCAGGTCGAAGGTCAAGATCTGCTCTCCAGGACTGTAGTCCAGCACGATCTGCCTCCCTATTGTGGTGCTGTATGCCAGTGGACAGCTCTTCGCCCACAGACAAGCGTAGCTGGACTCGAGCAGCTCACGCACTGTACTCGACCATTCCGTGCCGTCCGATGGACTCAAGCAGTCCTGCAGCAGCCAAGTTGCATCAACGCTGCTGTCGATATCGTCCTTCTCCAGCATCGTAGCTATCAGTGTCAAGTCTGCCTCAAGCAGAACACCCATACACGCTGAGAACGCTCGGCACTCGGGCGAGCTGCTGCAATCCTCCAATAACGTCGTGCACTCGCTGCAAGTTGGGGCTACATCATCCAGTTGTGAACCTGAATACAGTCCACCAAGTCGAGGCTGCTCCAAATCCAATGTTATGGTGATACCAGTGACTTCGTCTCCATCGTCGATCAGATCAGGAATCTCAGAGGTGATAAATGCTGGTTCAGGCTGGCCAGTGGTGTCTACCACCGCGAGGTCGCCAAACTGCCGACGTCGCACAATGGGGATGACAGTTGGGTCGTCAGGAACAAACTCCAGGTCTATGGTGAAGGGTGTGGTAGTGGTCATATCTTGCATCGAGTTTACAGCTTCGAACGTGAGACGGAAGCTCTGGGCATTGCCGCTTGATGACGACCCGCCTACGAAGAAGGACCACGAAGTATCCCCACCGAGACTCTGAGCACCAACCAGCACCCACGGAGCCGGATAAGTAGAAGTTGAACCAGCAACACTTGATCCGGGAGGAAGACCAATTGACGAAGATAGTGCAGCGTGAATGCTCTGCAGTTTACGAAGAAATATTGGAGCCAACTGATCGGCGCCTGTGAACCGGAGCTTGATACACAGTCTAGAAGTGGTGGAAGAGCACACGTCATCAGCATCTACCAATGCTCCAGATGATGCCGTCGACTCAGGGTTATCGAAGCCGTAGATAAAACTTGTTTGATGTTCCAGCAATGATTCCGACGAGGTAATGAAGCCGTCGCTGACCATGTCTATCTCTGGTACGGGATCTAACAGTCCAGGCAAATATTGCGGCCAAACGGGTGATCCATCGGCGTGGACGTCCATCTCTCGCTCAATCAGCTCGAGGCTGCCGAGTGAATACAAAGAAATCGTGGAAGCGAGTTGCTGCGAGGAAATGACGGAACGAATTCTTGCCATTGTTATGGTTGGTTTACGATAAATAAAGAATGATTGTCAGCTGGCTTAAACATAGAAATTTCTACCATGTTAAGTACACACAGTTGCCTCAAAATAGTCACGGTTTTAGCAAGAATCGAAATATTGACAAAAAGGGATCACTCCATACGACTATAGCGGTCATCAACCAAAGAGAAACAATTCTAAGTAGCGTGGGGGATGGTTTCTTTGCATTAGAAGTCACGTAGTTTTGTTTCAAGCCGCCATCAGTCTACTCTACTTTGGTCTTTATTCTCTGTTATAAAATAAAGCTGTCACTGCGCTTGTTCAGCGTTATATTTCATTTGTAAAGCGCCATCACCGATTCGTCACACTCGAATACCAAATGCTAGTATAAAAAAAAATTGTTCTACAAACGGCCTTCTTTTGAAAACAATCGTTCTTTCACAAAAAAAAACAACGAAAATGGAAAAAAGGAAATTGTATTAAATGCATAGACACAAAATCTGCAACTTCGCACTCAGTGCACACAGCTCGACCTTCGGCTTACAAGAAGTAGGCACCGGTACCGGGCCCTCACTTGTAGAGTTCATCCATGCCATGCGTGATCCCAGCAGCCGTCACGAACTCCAGCAGGACCATGTGGTCCCTCTTCTCGTTGGGGTCCTTGGAGAGGGCAGATTGCGTGGACAGGTAATGGTTGTCCGGCAGCAGGACAGGGCCATCGCCGATGGGCGTGTTCTGCTGGTAGTGGTCCGCCAGTTGCACGCTTCCATCTTCGATGTTGTGCCTGATCTTGAAGTTCACCTTGATGCCGTTCTTCTGCTTGTCGGCCATGATGTATACGTTGTGGGAGTTGTAGTTGTACTCCAGCTTGTGTCCGAGGATGTTTCCATCTTCCTTGAAATCGATGCCTTTCAGCTCGATGCGGTTCACCAGGGTATCACCTTCGAACTTGACTTCGGCACGTGTCTTGTAGTTCCCGTCATCCTTGAAGAAGATAGTCCTTTCTTGCACGTAGCCTTCGGGCATGGCGCTCTTGAAGAAGTCATGCTGCTTCATGTGATCTGGGTACCGGGAGAAGCACTGAACACCGTAGGTGAAAGTGGTGACGAGGGTTGGCCACGGAACAGGGAGCTTTCCGGTAGTGCAGATGAACTTCAGGGTGAGCTTTCCGTAGGTGGCATCACCTTCACCCTCTCCGCTGACGGAGAACTTGTGCCCGTTCACATCACCATCCAGTTCCACCAGGATTGGGACCACGCCAGTGAACAGTTCCTCGCCCTTGCCCATCTTAAGCTTGTAGAGTTCATCCATGCCATGCGTGATCCCAGCAGCCGTCACGAACTCCAGCAGGACCATGTGGTCCCTCTTCTCGTTGGGGTCCTTGGAGAGGGCAGATTGCGTGGACAGGTAATGGTTGTCCGGCAGCAGGACAGGGCCATCGCCGATGGGCGTGTTCTGCTGGTAGTGGTCCGCCAGTTGCACGCTTCCATCTTCGATGTTGTGCCTGATCTTGAAGTTCACCTTGATGCCGTTCTTCTGCTTGTCGGCCATGATGTATACGTTGTGGGAGTTGTAGTTGTACTCCAGCTTGTGTCCGAGGATGTTTCCATCTTCCTTGAAATCGATGCCTTTCAGCTCGATGCGGTTCACCAGGGTATCACCTTCGAACTTGACTTCGGCACGTGTCTTGTAGTTCCCGTCATCCTTGAAGAAGATAGTCCTTTCTTGCACGTAGCCTTCGGGCATGGCGCTCTTGAAGAAGTCATGCTGCTTCATGTGATCTGGGTACCGGGAGAAGCACTGAACACCGTAGGTGAAAGTGGTGACGAGGGTTGGCCACGGAACAGGGAGCTTTCCGGTAGTGCAGATGAACTTCAGGGTGAGCTTTCCGTAGGTGGCATCACCTTCACCCTCTCCGCTGACGGAGAACTTGTGCCCGTTCACATCACCATCCAGTTCCACCAGGATTGGGACCACGCCAGTGAACAGTTCCTCGCCCTTGCCCATACTAGTCCGCGGAGGCCTATCGATAAGCTTGATATCGAATTAATTCCTGCAGCCCGGGTTGTCGGTGAGAGTGAAGGTCGAAAAAGAGTCGGTTGGGACTTGGGCACGTGAGAGTGAAAAGGAGAATGAGCCTTCCGATCGTGGGCGAGTCGGGCGAGGCTTTTTGAGCGTCCGGTTGTGGCTTCACAACCCCACAAACAGGTGGTTTTCGGATGTAAAAGAGGAAGTGAAATGGCCAATCAGAAACGATCCCTGTGCAAGAGATGTCTTTAATGGGGCGCAAAAAAGACGATCAAAAAGAGCGGATAAGAGAGTTCCGGCGAGACATGTGGTGGTGGTGCTATAATTAACGGATGGTAGGACGCAAAAGCGCTCTGCACGGGTCAGGTTGAGAAGGTTCTCCAATCTTCTCTCTCTCTGTGACATGTGGTATTTCTCAGAAACCTTCGTGTAAACGCGAGGGAATGGACACGGAAATTAAGGTCTGAAAAAGGGTACTTGACTCGAGGTTTGTGGTTACGCCGGCTGAAGACGCCTAACCTCGCGTGCGATATGCCAGCTGTCGACATTGGAAAAGCCGAGATTGTCGCGGGTACGTAGAGGCGACCCTTTGTCCATCAGAGGGAGAGGAACCGACCAGTGGATAGGAGGGATAGACTTTATTTTGAGGGCGGTATAATGGTTATGGTAGCGACATTTTGATTTGACTTGAAAAAGATAAGAAAAAAAAAGGAAAAACTCGACTGCTTTGATGGGATAGCGATTTTTCCGTGGGTAGTCAGTTGGCCCTTGATTTTTACCATCAAGTACGTCCATATGACGGGAATAGGGACGCGGGTGGATATTAGGAGCGGCAGCTGACGTGGCAGCTCATAGACAAGGGGGTGCTTGAAAGGCTGTCTATGGTGGTGACGCCGTCTTGCGGTGTGGCAATGAGTTGTATCATTTAATCGAATACGAGGTATAA

**>pYF2.2-GFP**

AAGCTTGGCGTAATCATGGTCATAGCTGTTTCCTGTGTGAAATTGTTATCCGCTCACAATTCCACACAACATACGAGCCGGAAGCATAAAGTGTAAAGCCTGGGGTGCCTAATGAGTGAGCTAACTCACATTAATTGCGTTGCGCTCACTGCCCGCTTTCCAGTCGGGAAACCTGTCGTGCCAGCTGCATTAATGAATCGGCCAACGCGCGGGGAGAGGCGGTTTGCGTATTGGGCGCTCTTCCGCTTCCTCGCTCACTGACTCGCTGCGCTCGGTCGTTCGGCTGCGGCGAGCGGTATCAGCTCACTCAAAGGCGGTAATACGGTTATCCACAGAATCAGGGGATAACGCAGGAAAGAACATGTGAGCAAAAGGCCAGCAAAAGGCCAGGAACCGTAAAAAGGCCGCGTTGCTGGCGTTTTTCCATAGGCTCCGCCCCCCTGACGAGCATCACAAAAATCGACGCTCAAGTCAGAGGTGGCGAAACCCGACAGGACTATAAAGATACCAGGCGTTTCCCCCTGGAAGCTCCCTCGTGCGCTCTCCTGTTCCGACCCTGCCGCTTACCGGATACCTGTCCGCCTTTCTCCCTTCGGGAAGCGTGGCGCTTTCTCAATGCTCACGCTGTAGGTATCTCAGTTCGGTGTAGGTCGTTCGCTCCAAGCTGGGCTGTGTGCACGAACCCCCCGTTCAGCCCGACCGCTGCGCCTTATCCGGTAACTATCGTCTTGAGTCCAACCCGGTAAGACACGACTTATCGCCACTGGCAGCAGCCACTGGTAACAGGATTAGCAGAGCGAGGTATGTAGGCGGTGCTACAGAGTTCTTGAAGTGGTGGCCTAACTACGGCTACACTAGAAGGACAGTATTTGGTATCTGCGCTCTGCTGAAGCCAGTTACCTTCGGAAAAAGAGTTGGTAGCTCTTGATCCGGCAAACAAACCACCGCTGGTAGCGGTGGTTTTTTTGTTTGCAAGCAGCAGATTACGCGCAGAAAAAAAGGATCTCAAGAAGATCCTTTGATCTTTTCTACGGGGTCTGACGCTCAGTGGAACGAAAACTCACGTTAAGGGATTTTGGTCATGAGATTATCAAAAAGGATCTTCACCTAGATCCTTTTAAATTAAAAATGAAGTTTTAAATCAATCTAAAGTATATATGAGTAAACTTGGTCTGACAGTTACCAATGCTTAATCAGTGAGGCACCTATCTCAGCGATCTGTCTATTTCGTTCATCCATAGTTGCCTGACTCCCCGTCGTGTAGATAACTACGATACGGGAGGGCTTACCATCTGGCCCCAGTGCTGCAATGATACCGCGAGACCCACGCTCACCGGCTCCAGATTTATCAGCAATAAACCAGCCAGCCGGAAGGGCCGAGCGCAGAAGTGGTCCTGCAACTTTATCCGCCTCCATCCAGTCTATTAATTGTTGCCGGGAAGCTAGAGTAAGTAGTTCGCCAGTTAATAGTTTGCGCAACGTTGTTGCCATTGCTACAGGCATCGTGGTGTCACGCTCGTCGTTTGGTATGGCTTCATTCAGCTCCGGTTCCCAACGATCAAGGCGAGTTACATGATCCCCCATGTTGTGCAAAAAAGCGGTTAGCTCCTTCGGTCCTCCGATCGTTGTCAGAAGTAAGTTGGCCGCAGTGTTATCACTCATGGTTATGGCAGCACTGCATAATTCTCTTACTGTCATGCCATCCGTAAGATGCTTTTCTGTGACTGGTGAGTACTCAACCAAGTCATTCTGAGAATAGTGTATGCGGCGACCGAGTTGCTCTTGCCCGGCGTCAATACGGGATAATACCGCGCCACATAGCAGAACTTTAAAAGTGCTCATCATTGGAAAACGTTCTTCGGGGCGAAAACTCTCAAGGATCTTACCGCTGTTGAGATCCAGTTCGATGTAACCCACTCGTGCACCCAACTGATCTTCAGCATCTTTTACTTTCACCAGCGTTTCTGGGTGAGCAAAAACAGGAAGGCAAAATGCCGCAAAAAAGGGAATAAGGGCGACACGGAAATGTTGAATACTCATACTCTTCCTTTTTCAATATTATTGAAGCATTTATCAGGGTTATTGTCTCATGAGCGGATACATATTTGAATGTATTTAGAAAAATAAACAAATAGGGGTTCCGCGCACATTTCCCCGAAAAGTGCCACCTGACGTCTAAGAAACCATTATTATCATGACATTAACCTATAAAAATAGGCGTATCACGAGGCCCTTTCGTCTCGCGCGTTTCGGTGATGACGGTGAAAACCTCTGACACATGCAGCTCCCGGAGACGGTCACAGCTTGTCTGTAAGCGGATGCCGGGAGCAGACAAGCCCGTCAGGGCGCGTCAGCGGGTGTTGGCGGGTGTCGGGGCTGGCTTAACTATGCGGCATCAGAGCAGATTGTACTGAGAGTGCACCATATGCGGTGTGAAATACCGCACAGATGCGTAAGGAGAAAATACCGCATCAGGCGCCATTCGCCATTCAGGCTGCGCAACTGTTGGGAAGGGCGATCGGTGCGGGCCTCTTCGCTATTACGCCAGCTGGCGAAAGGGGGATGTGCTGCAAGGCGATTAAGTTGGGTAACGCCAGGGTTTTCCCAGTCACGACGTTGTAAAACGACGGCCAGTGAATTCCCCCAATTCCCCGGATCGTCTCATAGCTGCTCAAGACGTTTAGCAGATTGAGGATGCAGATCTTCTACTAATGAAACTACTGCAAACTACCAACAAAGAGGCGACCACAAAATGGACAGCCTATGCCGAGATTTGTTTAAGATACTCGATTTGCCGTCCAATTTTCATCAAGACGAAACACAGTGCATCGAACTGTGTTGTAAAAATTCATTTATTCTTGCAATTGTGACTCTCAAAGATTGATTAAACTTGAAAGCTGGTGAAAGCTTTCCTTCTTTTGGCATGTTTGTTACTAAGGCGGCGTAAGCACAATAGGCCCAGACTCATGCAAACCAGCCATAGTGACCTTCTCACTATGTTTCGCTTGAGATTTACTAGACTCTCGGGCATCTTCACAAAAGATGACAACCGAATCAACAAATTAAGTCAACGCTAAATCAAATATTTAGAAAATAAATTTTATTGTCGAAAAAACTCTGCTTTGACAATTTTCAAACTAACCAATTGCTAAGTATTCTAGTCGACACCGGTACCTATCAGAAGAACTCGTCAAGAAGGCGATAGAAGGCGATGCGCTGCGAATCGGGAGCGGCGATACCGTAAAGCACGAGGAAGCGGTCAGCCCATTCGCCGCCAAGCTCTTCAGCAATATCACGGGTAGCCAACGCTATGTCCTGATAGCGGTCCGCCACACCCAGCCGGCCACAGTCGATGAATCCAGAAAAGCGGCCATTTTCCACCATGATATTCGGCAAGCAGGCATCGCCATGGGTCACGACGAGATCCTCGCCGTCGGGCATGCGCGCCTTGAGCCTGGCGAACAGTTCGGCTGGCGCGAGCCCCTGATGCTCTTCGTCCAGATCATCCTGATCGACAAGACCGGCTTCCATCCGAGTACGTGCTCGCTCGATGCGATGTTTCGCTTGGTGGTCGAATGGGCAGGTAGCCGGATCAAGCGTATGCAGCCGCCGCATTGCATCAGCCATGATGGATACTTTCTCGGCAGGAGCAAGGTGAGATGACAGGAGATCCTGCCCCGGCACTTCGCCCAATAGCAGCCAGTCCCTTCCCGCTTCAGTGACAACGTCGAGCACAGCTGCGCAAGGAACGCCCGTCGTGGCCAGCCACGATAGCCGCGCTGCCTCGTCCTGCAGTTCATTCAGGGCACCGGACAGGTCGGTCTTGACAAAAAGAACCGGGCGCCCCTGCGCTGACAGCCGGAACACGGCGGCATCAGAGCAGCCGATTGTCTGTTGTGCCCAGTCATAGCCGAATAGCCTCTCCACCCAAGCGGCCGGAGAACCTGCGTGCAATCCATCTTGTTCAATCATTGTTGCCCGGGTGGATGCTCAGATGCTAGCGTCACCGAACGAATTAAAACTATAGTAAGTAAAGACTGTCGATGTAGCCATGTGGGGCCAATAGCTGCGAGGCTCAGAGAGAGACAAGGGGGTCGCCACTTACCCCGAACGATGTTCCAAGCAGAGCAGTCAGCAGAAAGTGAGGCTTGTTGCTGTCGGCGAGCGCAGCCTTGGGATCAACATGCGAGAGGATAGTTGGCTGCGAGCTTGCCAGCGTGGAGAGACTTGGTTATTGTAGGCAACAAGTCCCGCATGAAGGGCGTTGTATGGTGCAAAAAAGAGGAGATGGAATCTTCAAACACTTTGATTGGCTGGACAGCGGGCGCGGACTACTGGCCAACCAAAATGTGACCCCACTTGAGTGACACTTTCCCCACTATAGTGTCTTCAACTTGCACTGCAAGTGAAGTTGTTCTTGTTATTGCACTCGTTCTTACGTTATGGCTAGTACGCGCAGATGCAGAGCCTCCTCGGTGGTATTGCACGCCAACGAACCTCCATTCTCTGCGACATAAGGCTGTGGCAGTGCGCCCAGGAACTCGTAGAGAATGGTCAACTGCGCCGTCACCACTCCCGTGTTCGAGTCTTCCGTCGTGACGAGGGCCGACTTCACCGTCGCCACATTCGACAGCGCAGTGCGGTACATCTCCATCAGCATCGTGTCCAGTTGGGCTGTCACTTCCGTCGCGTTCGCCGTGAAGTCCTCGACAAAACTGTACGCCCAGTTGCCGAGCTCGAAGCTGACGCTGGCGAATCCGCTCGCCAGGTCGAAGGTCAAGATCTGCTCTCCAGGACTGTAGTCCAGCACGATCTGCCTCCCTATTGTGGTGCTGTATGCCAGTGGACAGCTCTTCGCCCACAGACAAGCGTAGCTGGACTCGAGCAGCTCACGCACTGTACTCGACCATTCCGTGCCGTCCGATGGACTCAAGCAGTCCTGCAGCAGCCAAGTTGCATCAACGCTGCTGTCGATATCGTCCTTCTCCAGCATCGTAGCTATCAGTGTCAAGTCTGCCTCAAGCAGAACACCCATACACGCTGAGAACGCTCGGCACTCGGGCGAGCTGCTGCAATCCTCCAATAACGTCGTGCACTCGCTGCAAGTTGGGGCTACATCATCCAGTTGTGAACCTGAATACAGTCCACCAAGTCGAGGCTGCTCCAAATCCAATGTTATGGTGATACCAGTGACTTCGTCTCCATCGTCGATCAGATCAGGAATCTCAGAGGTGATAAATGCTGGTTCAGGCTGGCCAGTGGTGTCTACCACCGCGAGGTCGCCAAACTGCCGACGTCGCACAATGGGGATGACAGTTGGGTCGTCAGGAACAAACTCCAGGTCTATGGTGAAGGGTGTGGTAGTGGTCATATCTTGCATCGAGTTTACAGCTTCGAACGTGAGACGGAAGCTCTGGGCATTGCCGCTTGATGACGACCCGCCTACGAAGAAGGACCACGAAGTATCCCCACCGAGACTCTGAGCACCAACCAGCACCCACGGAGCCGGATAAGTAGAAGTTGAACCAGCAACACTTGATCCGGGAGGAAGACCAATTGACGAAGATAGTGCAGCGTGAATGCTCTGCAGTTTACGAAGAAATATTGGAGCCAACTGATCGGCGCCTGTGAACCGGAGCTTGATACACAGTCTAGAAGTGGTGGAAGAGCACACGTCATCAGCATCTACCAATGCTCCAGATGATGCCGTCGACTCAGGGTTATCGAAGCCGTAGATAAAACTTGTTTGATGTTCCAGCAATGATTCCGACGAGGTAATGAAGCCGTCGCTGACCATGTCTATCTCTGGTACGGGATCTAACAGTCCAGGCAAATATTGCGGCCAAACGGGTGATCCATCGGCGTGGACGTCCATCTCTCGCTCAATCAGCTCGAGGCTGCCGAGTGAATACAAAGAAATCGTGGAAGCGAGTTGCTGCGAGGAAATGACGGAACGAATTCTTGCCATTGTTATGGTTGGTTTACGATAAATAAAGAATGATTGTCAGCTGGCTTAAACATAGAAATTTCTACCATGTTAAGTACACACAGTTGCCTCAAAATAGTCACGGTTTTAGCAAGAATCGAAATATTGACAAAAAGGGATCACTCCATACGACTATAGCGGTCATCAACCAAAGAGAAACAATTCTAAGTAGCGTGGGGGATGGTTTCTTTGCATTAGAAGTCACGTAGTTTTGTTTCAAGCCGCCATCAGTCTACTCTACTTTGGTCTTTATTCTCTGTTATAAAATAAAGCTGTCACTGCGCTTGTTCAGCGTTATATTTCATTTGTAAAGCGCCATCACCGATTCGTCACACTCGAATACCAAATGCTAGTATAAAAAAAAATTGTTCTACAAACGGCCTTCTTTTGAAAACAATCGTTCTTTCACAAAAAAAAACAACGAAAATGGAAAAAAGGAAATTGTATTAAATGCATAGACACAAAATCTGCAACTTCGCACTCAGTGCACACAGCTCGACCTTCGGCTTACAAGAAGTAGGCTCCGGAACCGGGCCCTCACTTGTAGAGTTCATCCATGCCATGCGTGATCCCAGCAGCCGTCACGAACTCCAGCAGGACCATGTGGTCCCTCTTCTCGTTGGGGTCCTTGGAGAGGGCAGATTGCGTGGACAGGTAATGGTTGTCCGGCAGCAGGACAGGGCCATCGCCGATGGGCGTGTTCTGCTGGTAGTGGTCCGCCAGTTGCACGCTTCCATCTTCGATGTTGTGCCTGATCTTGAAGTTCACCTTGATGCCGTTCTTCTGCTTGTCGGCCATGATGTATACGTTGTGGGAGTTGTAGTTGTACTCCAGCTTGTGTCCGAGGATGTTTCCATCTTCCTTGAAATCGATGCCTTTCAGCTCGATGCGGTTCACCAGGGTATCACCTTCGAACTTGACTTCGGCACGTGTCTTGTAGTTCCCGTCATCCTTGAAGAAGATAGTCCTTTCTTGCACGTAGCCTTCGGGCATGGCGCTCTTGAAGAAGTCATGCTGCTTCATGTGATCTGGGTACCGGGAGAAGCACTGAACACCGTAGGTGAAAGTGGTGACGAGGGTTGGCCACGGAACAGGGAGCTTTCCGGTAGTGCAGATGAACTTCAGGGTGAGCTTTCCGTAGGTGGCATCACCTTCACCCTCTCCGCTGACGGAGAACTTGTGCCCGTTCACATCACCATCCAGTTCCACCAGGATTGGGACCACGCCAGTGAACAGTTCCTCGCCCTTGCCCATCTTAAGCGTACGACTAGTCCGCGGAGGCCTATCGATAAGCTTGATATCGAATTAATTCCTGCAGCCCGGGTTGTCGGTGAGAGTGAAGGTCGAAAAAGAGTCGGTTGGGACTTGGGCACGTGAGAGTGAAAAGGAGAATGAGCCTTCCGATCGTGGGCGAGTCGGGCGAGGCTTTTTGAGCGTCCGGTTGTGGCTTCACAACCCCACAAACAGGTGGTTTTCGGATGTAAAAGAGGAAGTGAAATGGCCAATCAGAAACGATCCCTGTGCAAGAGATGTCTTTAATGGGGCGCAAAAAAGACGATCAAAAAGAGCGGATAAGAGAGTTCCGGCGAGACATGTGGTGGTGGTGCTATAATTAACGGATGGTAGGACGCAAAAGCGCTCTGCACGGGTCAGGTTGAGAAGGTTCTCCAATCTTCTCTCTCTCTGTGACATGTGGTATTTCTCAGAAACCTTCGTGTAAACGCGAGGGAATGGACACGGAAATTAAGGTCTGAAAAAGGGTACTTGACTCGAGGTTTGTGGTTACGCCGGCTGAAGACGCCTAACCTCGCGTGCGATATGCCAGCTGTCGACATTGGAAAAGCCGAGATTGTCGCGGGTACGTAGAGGCGACCCTTTGTCCATCAGAGGGAGAGGAACCGACCAGTGGATAGGAGGGATAGACTTTATTTTGAGGGCGGTATAATGGTTATGGTAGCGACATTTTGATTTGACTTGAAAAAGATAAGAAAAAAAAAGGAAAAACTCGACTGCTTTGATGGGATAGCGATTTTTCCGTGGGTAGTCAGTTGGCCCTTGATTTTTACCATCAAGTACGTCCATATGACGGGAATAGGGACGCGGGTGGATATTAGGAGCGGCAGCTGACGTGGCAGCTCATAGACAAGGGGGTGCTTGAAAGGCTGTCTATGGTGGTGACGCCGTCTTGCGGTGTGGCAATGAGTTGTATCATTTAATCGAATACGAGGTATAA

### Supplemental methods

#### Generation of *P. sojae* CRISPR/Cas9 plasmids

To express *hSpCas9* and sgRNAs effectively in *P. sojae*, we first created a new *Phytophthora* expression plasmid backbone pYF2 by combining elements from pHamT34 (Judelson *et al.*, 1991), pUN (Dou *et al.*, 2008) and pGFPN (Ah-Fong & Judelson, 2011) as follows. (i) The *HSP70* terminator was PCR amplified from pGFPN by using primers BlHSP70T_AgeI_F and BlHSP70T_NsiI_EcoRI_R which added an *Eco* RI site, and was then inserted into pUN using *Kpn* I and *Nsi* I sites, placing the *NPT II* gene under the control of the *RPL41* promoter and *HSP70* terminator. (ii) The entire *NPT II* cassette from (i) was then extracted by digestion with *Eco* RI and inserted into the *Eco* RI site of pHAMT34, resulting in pYF1. (iii) A synthetic multiple cloning site fragment containing the restriction enzyme sites *Xma I-Cla* I-*Stu* I-*Sac* II-*Spe* I-*Bsi* WI-*Afl* II-*Kpn* I was introduced between the *Xma* I and *Kpn* I sites of pYF1 by oligo annealing, creating pYF2. The plasmid pYF2-GFP was generated by adding an *eGFP* fragment amplified from pGFPN using primers GFP_AflII_F and GFP_ApaI_R into the restriction enzyme sites *Afl* II and *Apa* I of pYF2. The PsNLS, reported in Fang & Tyler (2015), was tested using the plasmid pYF2-PsNLS-2XGFP, which was constructed by two steps. (i) An extra *GFP* was amplified from the same plasmid pGFPN using primers GFP_SpeI_F and GFP_Afl II_R and inserted into the restriction sites *Spe* I and *Afl* II of the plasmid pYF2-GFP, generating pYF2-2XGFP. (ii) The PsNLS was inserted by annealing of two oligonucleotides encoding the NLS (MHKRKREDDTKVRRRMHKRKREDDTKVRRRMHKRKREDDTKVRRR).

To generate the construct for replacing the entire ORF of *Avr4/6*, the *NPT II* coding region together with 250 bp, 500 bp or 1 kb of 5**’** and 3**’** flanking regions outside the *Avr4/6* coding region were PCR amplified and cloned into the plasmid pBluescript II KS+ by In-Fusion® HD Cloning Kit (Clontech).

### Supplemental Sequences

#### DNA template used for CRISPR *in vitro* cleavage assay

The partial Avr4/6 sequence (in red) was TA-cloned in pCR2 (Invitrogen). Targets of sgRNA-A and sgRNA-B are highlighted in blue and yellow respectively. Cleavage sites are indicated by arrows. PCR product (454bp) amplified by M13F/M13R was used as the DNA template for CRISPR *in vitro* cleavage assay. Expected fragment sizes (bp) after cleaved by Cas9, sgRNA-A: 136 and 318; sgRNA-B: 269 and 185.

>pCR2_Avr4/6 partial

M13R <u>CAGGAAACAGCTATGAC</u>CATGATTACGCCAAGCTTGGTACCGAGCTCGGATCCACT AGTAACGGCCGCCAGTGTGCTGGAATTCGGCTTTTGTTGCCGCTCCAGCTGATGCGATCACAGATGAGTCTCAGCCCCGC↓GATGCAACCATCGTCGATGCCCCACTCACTGGCAGGGGTGCCAATGCTCGGTATTTACGGACTAGCACATCGATCATCAAGGCCCCCGACGCCCAGCTACCGAGTACAAAGGCTGCGATCGCATCGTCTGTCA↓CCAAGG AAGAAGAGGAACGCAAGATCTCGACCGGTCTCAGCAAGCTCAGGCAGAAGCTGAGCAAGCGTTTTCACAAGCCGAATTCTGCAGATATCCATCACACTGGCGGCCGCTCGA GCATGCATCTAGAGGGCCCAATTCGCCCTATAGTGAGTCGTATTACAATTCA<u>CTGG CCGTCGTTTTAC</u>

M13F

#### Sequences of sgRNA and ribozyme flanked sgRNAs

Nucleotides highlighted in pink: the 20nt target sequence

Nucleotides highlighted in yellow: 80nt sgRNA scaffold

Nucleotides highlighted in green: hammerhead ribozymes (HH ribozymes)

Nucleotides highlighted in cyan: HDV ribozymes

<u>Nucleotides underline</u>: The first six nucleotides of the Hammerhead (HH) ribozyme must be complementary to the first six nucleotides of the target sequence.

**↓**: ribozyme cleavage sites

#### >Target A without Ribozyme

TCGACGATGGTTGCATCGCGgttttagagctagaaatagcaagttaaaataaggctagtccgttatcaacttgaaaaagtggcaccgagtcggtgctttt

#### >Target B without Ribozyme

GATCGCATCGTCTGTCACCAgttttagagctagaaatagcaagttaaaataaggctagtccgttatcaacttgaaaaagtggcaccgagtcggtgctttt

#### >Target A with Ribozyme

<u>cgtcga</u>**ctgatgagtccgtgaggacgaaacgagtaagctcgtc↓**<u>TCGACG</u>ATGGTTGCATCGCGgtttt agagctagaaatagcaagttaaaataaggctagtccgttatcaacttgaaaaagtggcaccgagtcggtgctttt**↓ggccg gcatggtcccagcctcctcgctggcgccggctgggcaacatgcttcggcatggcgaatgggac**

#### >Target B with Ribozyme

<u>gcgatc</u>**ctgatgagtccgtgaggacgaaacgagtaagctcgtc↓**<u>GATCGC</u>ATCGTCTGTCACCAgtttta gagctagaaatagcaagttaaaataaggctagtccgttatcaacttgaaaaagtggcaccgagtcggtgctttt**↓ggccgg catggtcccagcctcctcgctggcgccggctgggcaacatgcttcggcatggcgaatgggac**

#### >PsNLS-Cas9

*P. sojae* NLS in red, Restriction enzymes used for cloning are underscored. Start and stop codon are in bold.

**ATG**CACAAGCGCAAGCGCGAGGACGACACCAAGGTCCGTCGTCGCATGCACAAGCGCAAGCGCGAGGACGACACCAAGGTCCGTCGTCGCATGCACAAGCGCAAGCGCGAGGACGACACCAAGGTCCGTCGTCGC<u>ACTAGT</u>ATGGCCCCAAAGAAGAAGCGGAAGGTCGGTATCCACGGAGTCCCAGCAGCCGACAAGAAGTACAGCATCGGCCTGGACATCGGCACCAACTCTGTGGGCTGGGCCGTGATCACCGACGAGTACAAGGTGCCCAGCAAGAAATTCAAGGTGCTGGGCAACACCGACCGGCACAGCATCAAGAAGAACCTGATCGGAGCCCTGCTGTTCGACAGCGGCGAAACAGCCGAGGCCACCCGGCTGAAGAGAACCGCCAGAAGAAGATACACCAGACGGAAGAACCGGATCTGCTATCTGCAAGAGATCTTCAGCAACGAGATGGCCAAGGTGGACGACAGCTTCTTCCACAGACTGGAAGAGTCCTTCCTGGTGGAAGAGGATAAGAAGCACGAGCGGCACCCCATCTTCGGCAACATCGTGGACGAGGTGGCCTACCACGAGAAGTACCCCACCATCTACCACCTGAGAAAGAAACTGGTGGACAGCACCGACAAGGCCGACCTGCGGCTGATCTATCTGGCCCTGGCCCACATGATCAAGTTCCGGGGCCACTTCCTGATCGAGGGCGACCTGAACCCCGACAACAGCGACGTGGACAAGCTGTTCATCCAGCTGGTGCAGACCTACAACCAGCTGTTCGAGGAAAACCCCATCAACGCCAGCGGCGTGGACGCCAAGGCCATCCTGTCTGCCAGACTGAGCAAGAGCAGACGGCTGGAAAATCTGATCGCCCAGCTGCCCGGCGAGAAGAAGAATGGCCTGTTCGGAAACCTGATTGCCCTGAGCCTGGGCCTGACCCCCAACTTCAAGAGCAACTTCGACCTGGCCGAGGATGCCAAACTGCAGCTGAGCAAGGACACCTACGACGACGACCTGGACAACCTGCTGGCCCAGATCGGCGACCAGTACGCCGACCTGTTTCTGGCCGCCAAGAACCTGTCCGACGCCATCCTGCTGAGCGACATCCTGAGAGTGAACACCGAGATCACCAAGGCCCCCCTGAGCGCCTCTATGATCAAGAGATACGACGAGCACCACCAGGACCTGACCCTGCTGAAAGCTCTCGTGCGGCAGCAGCTGCCTGAGAAGTACAAAGAGATTTTCTTCGACCAGAGCAAGAACGGCTACGCCGGCTACATTGACGGCGGAGCCAGCCAGGAAGAGTTCTACAAGTTCATCAAGCCCATCCTGGAAAAGATGGACGGCACCGAGGAACTGCTCGTGAAGCTGAACAGAGAGGACCTGCTGCGGAAGCAGCGGACCTTCGACAACGGCAGCATCCCCCACCAGATCCACCTGGGAGAGCTGCACGCCATTCTGCGGCGGCAGGAAGATTTTTACCCATTCCTGAAGGACAACCGGGAAAAGATCGAGAAGATCCTGACCTTCCGCATCCCCTACTACGTGGGCCCTCTGGCCAGGGGAAACAGCAGATTCGCCTGGATGACCAGAAAGAGCGAGGAAACCATCACCCCCTGGAACTTCGAGGAAGTGGTGGACAAGGGCGCTTCCGCCCAGAGCTTCATCGAGCGGATGACCAACTTCGATAAGAACCTGCCCAACGAGAAGGTGCTGCCCAAGCACAGCCTGCTGTACGAGTACTTCACCGTGTATAACGAGCTGACCAAAGTGAAATACGTGACCGAGGGAATGAGAAAGCCCGCCTTCCTGAGCGGCGAGCAGAAAAAGGCCATCGTGGACCTGCTGTTCAAGACCAACCGGAAAGTGACCGTGAAGCAGCTGAAAGAGGACTACTTCAAGAAAATCGAGTGCTTCGACTCCGTGGAAATCTCCGGCGTGGAAGATCGGTTCAACGCCTCCCTGGGCACATACCACGATCTGCTGAAAATTATCAAGGACAAGGACTTCCTGGACAATGAGGAAAACGAGGACATTCTGGAAGATATCGTGCTGACCCTGACACTGTTTGAGGACAGAGAGATGATCGAGGAACGGCTGAAAACCTATGCCCACCTGTTCGACGACAAAGTGATGAAGCAGCTGAAGCGGCGGAGATACACCGGCTGGGGCAGGCTGAGCCGGAAGCTGATCAACGGCATCCGGGACAAGCAGTCCGGCAAGACAATCCTGGATTTCCTGAAGTCCGACGGCTTCGCCAACAGAAACTTCATGCAGCTGATCCACGACGACAGCCTGACCTTTAAAGAGGACATCCAGAAAGCCCAGGTGTCCGGCCAGGGCGATAGCCTGCACGAGCACATTGCCAATCTGGCCGGCAGCCCCGCCATTAAGAAGGGCATCCTGCAGACAGTGAAGGTGGTGGACGAGCTCGTGAAAGTGATGGGCCGGCACAAGCCCGAGAACATCGTGATCGAAATGGCCAGAGAGAACCAGACCACCCAGAAGGGACAGAAGAACAGCCGCGAGAGAATGAAGCGGATCGAAGAGGGCATCAAAGAGCTGGGCAGCCAGATCCTGAAAGAACACCCCGTGGAAAACACCCAGCTGCAGAACGAGAAGCTGTACCTGTACTACCTGCAGAATGGGCGGGATATGTACGTGGACCAGGAACTGGACATCAACCGGCTGTCCGACTACGATGTGGACCATATCGTGCCTCAGAGCTTTCTGAAGGACGACTCCATCGACAACAAGGTGCTGACCAGAAGCGACAAGAACCGGGGCAAGAGCGACAACGTGCCCTCCGAAGAGGTCGTGAAGAAGATGAAGAACTACTGGCGGCAGCTGCTGAACGCCAAGCTGATTACCCAGAGAAAGTTCGACAATCTGACCAAGGCCGAGAGAGGCGGCCTGAGCGAACTGGATAAGGCCGGCTTCATCAAGAGACAGCTGGTGGAAACCCGGCAGATCACAAAGCACGTGGCACAGATCCTGGACTCCCGGATGAACACTAAGTACGACGAGAATGACAAGCTGATCCGGGAAGTGAAAGTGATCACCCTGAAGTCCAAGCTGGTGTCCGATTTCCGGAAGGATTTCCAGTTTTACAAAGTGCGCGAGATCAACAACTACCACCACGCCCACGACGCCTACCTGAACGCCGTCGTGGGAACCGCCCTGATCAAAAAGTACCCTAAGCTGGAAAGCGAGTTCGTGTACGGCGACTACAAGGTGTACGACGTGCGGAAGATGATCGCCAAGAGCGAGCAGGAAATCGGCAAGGCTACCGCCAAGTACTTCTTCTACAGCAACATCATGAACTTTTTCAAGACCGAGATTACCCTGGCCAACGGCGAGATCCGGAAGCGGCCTCTGATCGAGACAAACGGCGAAACCGGGGAGATCGTGTGGGATAAGGGCCGGGATTTTGCCACCGTGCGGAAAGTGCTGAGCATGCCCCAAGTGAATATCGTGAAAAAGACCGAGGTGCAGACAGGCGGCTTCAGCAAAGAGTCTATCCTGCCCAAGAGGAACAGCGATAAGCTGATCGCCAGAAAGAAGGACTGGGACCCTAAGAAGTACGGCGGCTTCGACAGCCCCACCGTGGCCTATTCTGTGCTGGTGGTGGCCAAAGTGGAAAAGGGCAAGTCCAAGAAACTGAAGAGTGTGAAAGAGCTGCTGGGGATCACCATCATGGAAAGAAGCAGCTTCGAGAAGAATCCCATCGACTTTCTGGAAGCCAAGGGCTACAAAGAAGTGAAAAAGGACCTGATCATCAAGCTGCCTAAGTACTCCCTGTTCGAGCTGGAAAACGGCCGGAAGAGAATGCTGGCCTCTGCCGGCGAACTGCAGAAGGGAAACGAACTGGCCCTGCCCTCCAAATATGTGAACTTCCTGTACCTGGCCAGCCACTATGAGAAGCTGAAGGGCTCCCCCGAGGATAATGAGCAGAAACAGCTGTTTGTGGAACAGCACAAGCACTACCTGGACGAGATCATCGAGCAGATCAGCGAGTTCTCCAAGAGAGTGATCCTGGCCGACGCTAATCTGGACAAAGTGCTGTCCGCCTACAACAAGCACCGGGATAAGCCCATCAGAGAGCAGGCCGAGAATATCATCCACCTGTTTACCCTGACCAATCTGGGAGCCCCTGCCGCCTTCAAGTACTTTGACACCACCATCGACCGGAAGAGGTACACCAGCACCAAAGAGGTGCTGGACGCCACCCTGATCCACCAGAGCATCACCGGCCTGTACGAGACACGGATCGACCTGTCTCAGCTGGGAGGCGAC<u>CTTAAG</u>CCG<u>GGGCCC</u>**TAA**

### Sequence of plasmid having donor DNA

*NPT II* gene is in gray background. Nucleotides in green, pink are the border of, 250 bp, 500 bp and 1 kb homologous arms.

>pBS_KS_Avr4/6-1k

CTAAATTGTAAGCGTTAATATTTTGTTAAAATTCGCGTTAAATTTTTGTTAAATCAGCTCATTTTTTAACCAATAGGCCGAAATCGGCAAAATCCCTTATAAATCAAAAGAATAGACCGAGATAGGGTTGAGTGTTGTTCCAGTTTGGAACAAGAGTCCACTATTAAAGAACGTGGACTCCAACGTCAAAGGGCGAAAAACCGTCTATCAGGGCGATGGCCCACTACGTGAACCATCACCCTAATCAAGTTTTTTGGGGTCGAGGTGCCGTAAAGCACTAAATCGGAACCCTAAAGGGAGCCCCCGATTTAGAGCTTGACGGGGAAAGCCGGCGAACGTGGCGAGAAAGGAAGGGAAGAAAGCGAAAGGAGCGGGCGCTAGGGCGCTGGCAAGTGTAGCGGTCACGCTGCGCGTAACCACCACACCCGCCGCGCTTAATGCGCCGCTACAGGGCGCGTCCCATTCGCCATTCAGGCTGCGCAACTGTTGGGAAGGGCGATCGGTGCGGGCCTCTTCGCTATTACGCCAGCTGGCGAAAGGGGGATGTGCTGCAAGGCGATTAAGTTGGGTAACGCCAGGGTTTTCCCAGTCACGACGTTGTAAAACGACGGCCAGTGAGCGCGCGTAATACGACTCACTATAGGGCGAATTGGGTACCGGGCCCCCCCTCGAGGTCGACGGTATCGATAAGCTTGATATCGAATTCGTAGGAAATCTCAAATTCGGTATACCTACCATATTTAGTTATAATGGGGGTGGTACAAAATGGGAATGAGTACTTTCCTACCTCGATCATCAATCACTTCATACAAACTTTTCATAAAGACTTCGCCCGTCTCCAAAATGGCTCCTTTTTCGGCAACCCCATTATAACTAAACCCCCACAACAACTAAATAAATACGGTAAATGTTATAGGTGCCCTGCAGCTTGTCCGACATGTACATCGTTCATCATCCGTACGTTTTGGAGTTGGCTCGCCTGTTCTCATCAATGCCGTCGATCGTAGCAATTCCTTACTCCGTTTTCTGGTGTGTACAGTATCGGGTCGTAAACACATGCGTAAACACATGCGTAATGCGATACGGCGCGGTGTCTCAGAACCACCCGCCGATTTTGTACTGACAGCGTACCATTGACCTGTAGAGCCCACCCAGATCATAATTAATCTTCCAATAAGCAAGCGCCCTGCAATTCCAAGTCCAACGatgattgaacaagatggattgcacgcaggttctccggccgcttgggtggagaggctattcggctatgactgggcacaacagacaatcggctgctctgatgccgccgtgttccggctgtcagcgcaggggcgcccggttctttttgtcaagaccgacctgtccggtgccctgaatgaactgcaggacgaggcagcgcggctatcgtggctggccacgacgggcgttccttgcgcagctgtgctcgacgttgtcactgaagcgggaagggactggctgctattgggcgaagtgccggggcaggatctcctgtcatctcaccttgctcctgccgagaaagtatccatcatggctgatgcaatgcggcggctgcatacgcttgatccggctacctgcccattcgaccaccaagcgaaacatcgcatcgagcgagcacgtactcggatggaagccggtcttgtcgatcaggatgatctggacgaagagcatcaggggctcgcgccagccgaactgttcgccaggctcaaggcgcgcatgcccgacggcgaggatctcgtcgtgacccatggcgatgcctgcttgccgaatatcatggtggaaaatggccgcttttctggattcatcgactgtggccggctgggtgtggcggaccgctatcaggacatagcgttggctacccgtgatattgctgaagagcttggcggcgaatgggctgaccgcttcctcgtgctttacggtatcgccgctcccgattcgcagcgcatcgccttctatcgccttcttgacgagttcttctgaGTTTCACAGGCGTTCTGGCTGGAAATCCCGCAATCGTGCCAATACGGGAAAGCACTCGTATTTGTATCCATAAGCATATTCCTGACTCGATCAGTATCTGAGCCGCTAAAACAACTTCATAGCACCACATTTCTATAGAATTGTAAACCCTGAGAGCAAAGAATCGAACCTTCATTTAGCCTTTCTCTTTGAAAAAAGCGCCGGAGTCTGTGTGGCTCGTGCTGGGACACTGGAACCCACTAGAACAGGCTGATGAGCATAAGGTTCACGCCACTCAGTATGACCGAGATATCCGGTAAACCCACTTGCAGCTTTTTCAAAACCCTTTTTAAGGCGCTCGGGGACCCTTTCGGCCTCTGGAGCAGCTTGCCAAGACCGCTCGATATCCCTCCCTCTTCGTCGACGCCTAGCGAAGTGTGCATCAGCTGGCTGACGGATGCCTTGGTCGAGTCGTCACTCCGGAGTTGTCTCGCATCGGGACTAAGTAAGCGAGGCGCAGCGGATCCACTAGTTCTAGAGCGGCCGCCACCGCGGTGGAGCTCCAGCTTTTGTTCCCTTTAGTGAGGGTTAATTGCGCGCTTGGCGTAATCATGGTCATAGCTGTTTCCTGTGTGAAATTGTTATCCGCTCACAATTCCACACAACATACGAGCCGGAAGCATAAAGTGTAAAGCCTGGGGTGCCTAATGAGTGAGCTAACTCACATTAATTGCGTTGCGCTCACTGCCCGCTTTCCAGTCGGGAAACCTGTCGTGCCAGCTGCATTAATGAATCGGCCAACGCGCGGGGAGAGGCGGTTTGCGTATTGGGCGCTCTTCCGCTTCCTCGCTCACTGACTCGCTGCGCTCGGTCGTTCGGCTGCGGCGAGCGGTATCAGCTCACTCAAAGGCGGTAATACGGTTATCCACAGAATCAGGGGATAACGCAGGAAAGAACATGTGAGCAAAAGGCCAGCAAAAGGCCAGGAACCGTAAAAAGGCCGCGTTGCTGGCGTTTTTCCATAGGCTCCGCCCCCCTGACGAGCATCACAAAAATCGACGCTCAAGTCAGAGGTGGCGAAACCCGACAGGACTATAAAGATACCAGGCGTTTCCCCCTGGAAGCTCCCTCGTGCGCTCTCCTGTTCCGACCCTGCCGCTTACCGGATACCTGTCCGCCTTTCTCCCTTCGGGAAGCGTGGCGCTTTCTCATAGCTCACGCTGTAGGTATCTCAGTTCGGTGTAGGTCGTTCGCTCCAAGCTGGGCTGTGTGCACGAACCCCCCGTTCAGCCCGACCGCTGCGCCTTATCCGGTAACTATCGTCTTGAGTCCAACCCGGTAAGACACGACTTATCGCCACTGGCAGCAGCCACTGGTAACAGGATTAGCAGAGCGAGGTATGTAGGCGGTGCTACAGAGTTCTTGAAGTGGTGGCCTAACTACGGCTACACTAGAAGGACAGTATTTGGTATCTGCGCTCTGCTGAAGCCAGTTACCTTCGGAAAAAGAGTTGGTAGCTCTTGATCCGGCAAACAAACCACCGCTGGTAGCGGTGGTTTTTTTGTTTGCAAGCAGCAGATTACGCGCAGAAAAAAAGGATCTCAAGAAGATCCTTTGATCTTTTCTACGGGGTCTGACGCTCAGTGGAACGAAAACTCACGTTAAGGGATTTTGGTCATGAGATTATCAAAAAGGATCTTCACCTAGATCCTTTTAAATTAAAAATGAAGTTTTAAATCAATCTAAAGTATATATGAGTAAACTTGGTCTGACAGTTACCAATGCTTAATCAGTGAGGCACCTATCTCAGCGATCTGTCTATTTCGTTCATCCATAGTTGCCTGACTCCCCGTCGTGTAGATAACTACGATACGGGAGGGCTTACCATCTGGCCCCAGTGCTGCAATGATACCGCGAGACCCACGCTCACCGGCTCCAGATTTATCAGCAATAAACCAGCCAGCCGGAAGGGCCGAGCGCAGAAGTGGTCCTGCAACTTTATCCGCCTCCATCCAGTCTATTAATTGTTGCCGGGAAGCTAGAGTAAGTAGTTCGCCAGTTAATAGTTTGCGCAACGTTGTTGCCATTGCTACAGGCATCGTGGTGTCACGCTCGTCGTTTGGTATGGCTTCATTCAGCTCCGGTTCCCAACGATCAAGGCGAGTTACATGATCCCCCATGTTGTGCAAAAAAGCGGTTAGCTCCTTCGGTCCTCCGATCGTTGTCAGAAGTAAGTTGGCCGCAGTGTTATCACTCATGGTTATGGCAGCACTGCATAATTCTCTTACTGTCATGCCATCCGTAAGATGCTTTTCTGTGACTGGTGAGTACTCAACCAAGTCATTCTGAGAATAGTGTATGCGGCGACCGAGTTGCTCTTGCCCGGCGTCAATACGGGATAATACCGCGCCACATAGCAGAACTTTAAAAGTGCTCATCATTGGAAAACGTTCTTCGGGGCGAAAACTCTCAAGGATCTTACCGCTGTTGAGATCCAGTTCGATGTAACCCACTCGTGCACCCAACTGATCTTCAGCATCTTTTACTTTCACCAGCGTTTCTGGGTGAGCAAAAACAGGAAGGCAAAATGCCGCAAAAAAGGGAATAAGGGCGACACGGAAATGTTGAATACTCATACTCTTCCTTTTTCAATATTATTGAAGCATTTATCAGGGTTATTGTCTCATGAGCGGATACATATTTGAATGTATTTAGAAAAATAAACAAATAGGGGTTCCGCGCACATTTCCCCGAAAA GTGCCAC

